# Microtubule polymerization state and clathrin-dependent internalization regulate dynamics of cardiac potassium channel

**DOI:** 10.1101/805234

**Authors:** Dario Melgari, Camille Barbier, Gilles Dilanian, Catherine Rücker-Martin, Nicolas Doisne, Alain Coulombe, Stéphane N. Hatem, Elise Balse

**Affiliations:** INSERM UMRS1166, ICAN - Institute of CardioMetabolism and Nutrition, Sorbonne Université, Paris, France; Institut de Cardiologie, Hôpital Pitié-Salpêtrière, Paris, France; INSERM U999, University of Paris-Sud and Centre Chirurgical Marie Lannelongue, France

**Author notes:** Corresponding author: Dr. Elise Balse, PhD, INSERM UMRS1166, Faculté de Médecine Pitié-Salpêtrière, 91, Boulevard de l’Hôpital, 75013 Paris.

**Keywords:** atrial myocytes, potassium channel dynamics, clathrin, cytoskeleton, atrial remodeling

## Abstract

Ion channel trafficking powerfully influences cardiac electrical activity as it regulates the number of available channels at the plasma membrane. Studies have largely focused on identifying the molecular determinants of the trafficking of the atria-specific K_V_1.5 channel, the molecular basis of the ultra-rapid delayed rectifier current *I*_Kur_. Besides, regulated K_V_1.5 channel recycling upon changes in homeostatic state and mechanical constraints in native cardiomyocytes has been well documented. Here, using cutting-edge imaging in live myocytes, we investigated the dynamics of this channel in the plasma membrane. We demonstrate that the clathrin pathway is a major regulator of the functional expression of K_V_1.5 channels in atrial myocytes, with the microtubule network as the prominent organizer of K_V_1.5 transport within the membrane. Both clathrin blockade and microtubule disruption result in channel clusturization with reduced membrane mobility and internalization, whereas disassembly of the actin cytoskeleton does not. Mobile K_V_1.5 channels are associated with the microtubule plus-end tracking protein EB1 whereas static K_V_1.5 clusters are associated with stable acetylated microtubules. In human biopsies from patients in atrial fibrillation associated with atrial remodeling, drastic modifications in the trafficking balance occurs together with alteration in microtubule polymerization state resulting in modest reduced endocytosis and increased recycling. Consequently, hallmark of atrial K_V_1.5 dynamics within the membrane is clathrin- and microtubule-dependent. During atrial remodeling, predominance of anterograde trafficking activity over retrograde trafficking could result in accumulation ok K_V_1.5 channels in the plasma membrane.

## INTRODUCTION

The current understanding of basic cardiac electrophysiology has undergone a profound transformation. The classical theory that the cardiac action potential is generated by the successive opening or closing of different ion channels with distinct biophysical properties is now challenged by accumulating evidence indicating that these proteins belong to a large molecular network undergoing continuous dynamic regulation. Thus, functional ion channel expression results from highly-integrated cellular activities including post-translational regulation, trafficking, and cell microarchitecture organization.

The pathophysiology of cardiac arrhythmias has begun to be revisited in the light of this new paradigm. A first clue was the discovery of mutations in the *HERG* gene producing a trafficking-defective channel, and reduced corresponding *I*_Kr_ current amplitude, in inherited long-QT syndrome^1^. Another example is the observation that hypokalemia, a risk factor for cardiac arrhythmias, reduces hERG channel density at the plasma membrane by promoting channel internalization^2^. We previously reported that K_V_1.5 channels, the molecular basis of the atria-specific ultra-rapid delayed rectifier current *I*_Kur_, are stored in submembrane recycling endosomes and are ready for recruitment upon changes in membrane cholesterol content^3^ or upon shear stress^4^. The latter process requires activation of the integrin/phosphorylated FAK signaling pathway which becomes constitutively stimulated during atrial dilation as a result of excessive atrial wall cardiac myocyte shear stress, thereby contributing to action potential shortening and increased susceptibility to atrial arrhythmias.

The aim of the present study was to further decipher the trajectory and trafficking pathways of K_V_1.5 channels and the contribution of these processes to potassium current properties in atrial cardiomyocytes. Specifically, we investigated the role of the cytoskeleton and internalization routes involved in functional K_V_1.5 channel expression. There is currently limited information about K_V_1.5 channel internalization, reported to be regulated by both dynamin- and dynein-dependent pathways in heterologous expression systems^5,6^. It has been suggested that K_V_1.5 internalization could occur via cavaolae association^7,8^, however the clathrin pathway has not yet been investigated. Using whole cell patch clamp, biochemistry techniques, and cutting-edge imaging approaches, we showed that K_V_1.5 channels are highly mobile in the sarcolemma and follow the microtubule network, whereas clathrin pathway is involved as the principal internalization route. In remodeled atrial myocardium from patients in atrial fibrillation (AF), we provide evidence for reduced endocytosis and increased recycling that could contribute to the accumulation of K_V_1.5 channels in the sarcolemma of atrial myocytes during disease.

## RESULTS

### K_V_1.5 channels are associated with clathrin vesicles but not with caveolae

We first investigated the association of K_V_1.5 channels with canonical endocytosis pathways in freshly isolated atrial myocytes using high resolution 3-dimensional (3-D) deconvolution microscopy. Colocalization analysis revealed that K_V_1.5 channels co-localized poorly with caveolin 3 (R: 0.08) but associated with clathrin vesicles (R: 0.53) labelled with a clathrin light chain (LC) antibody (Figure 1A, C) at cell boundaries (n=5 cells, 15 lines/cell). Immunofluorescence images showed that clathrin vesicles were located at the intercalated disc (ID), the lateral membrane (LM), and as expected, at the perinuclear level (Figure S1, S2). K_V_1.5 and clathrin LC-positives vesicles were located both at the ID and at the LM (Figure 1C). Electron microscopy images also revealed the presence of clathrin coated vesicles (CCVs) and clathrin buds at the ID and at the LM (Figure S1B). Interestingly, CCVs were also observable along the z-lines in atrial sections. Quantification of the number of clathrin buds and CCVs at the ID and the LM showed that the ID displayed an equal amount of buds and internalized vesicles whereas the number of internalized vesicles was superior to the number of buds at the LM, suggesting that the LM is a more active endocytosis zone (Figure S1D). Immunostaining experiments were also performed in cultured atrial myocytes in which the human K_V_1.5 channel fused to the GFP protein (h-K_V_1.5-GFP) was overexpressed using an adenoviral vector. In overexpression condition as well, the association of K_V_1.5 was observed only with clathrin vesicles (Figure 1B, D). These observations show that clathrin-dependent endocytosis is likely to be the internalization pathway of K_V_1.5 channels in atrial myocytes.

**Figure 1.**
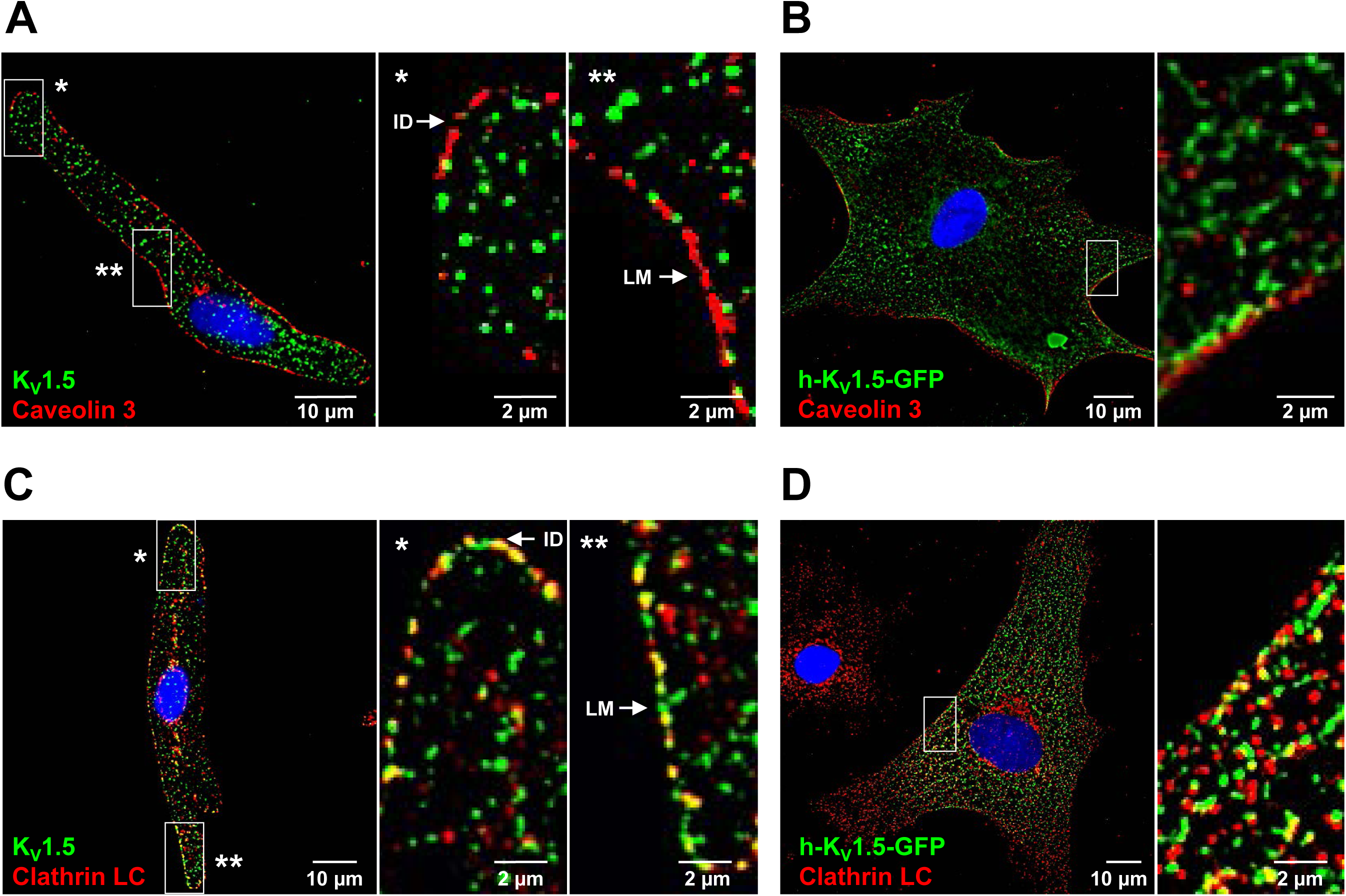
K_V_1.5 channels are associated with clathrin vesicles but not with caveolae in atrial myocytes. **A, C**, Example microscopy images of freshly isolated atrial myocytes stained with K_V_1.5 and (**A**) caveolin 3 or (**C**) clathrin light-chain (LC), and enlargements of two regions of interest (ROI * and **). **B, D**, Example microscopy images of cultured atrial cardiomyocytes expressing h-K_V_1.5-GFP and stained with (**B**) caveolin 3 or (**D**) clathrin-LC, and enlargement of one ROI. LM: Lateral Membrane; ID: Intercalated Disc.

### Clathrin pathway regulates the density of functional K_V_1.5 channels in atrial myocytes

Biotinylation experiments were performed to quantify the surface expression of h-K_V_1.5-GFP channels after blockade of the clathrin pathway with sucrose (225 mmol/L) treatment for 2 hours. Sucrose prevents clathrin from interacting with the adaptor protein complex AP2 and triggers abnormal formation of microcages^9^. The h-K_V_1.5-GFP band was ∼1.5-fold stronger in the membrane fraction after sucrose treatment (Figure 2A, n=3). The transferrin receptor was used as a control of clathrin-mediated endocytosis and was increased by 3 fold in the surface fraction (Figure 2A). Next, surface fluorescence signal was quantified in cultured atrial myocytes transduced with h-K_V_1.5-HA, the HA tag being located on K_V_1.5 extracellular loop. Cells were incubated for 2 hours with control medium or sucrose and live stained with anti-HA and secondary Fab A48 antibody before fixation allowing visualization of only pools of channels expressed at the surface. In line with biotinylation assays, quantification using high resolution 3-D deconvolution microscopy revealed ∼40% increase in K_V_1.5 surface staining in sucrose-treated cells at T0 (Figure 2B). Internalization assay further supported the importance of the clathrin pathway on K_V_1.5 channel endocytosis: whereas ∼40% of channels were internalized after 2 hours (T2) in control condition, internalization was almost completely prevented in sucrose-treated cells (15%, not significantly different from T0) (Figure 2B). Perspective views of 3D projection of 4 µm stack acquired at 0.2 µm intervals also showed the HA staining at the membrane at T0 and reduced internalization in sucrose-treated cells compared to control at T2 (Figure 2C). The effect of clathrin blockade on K_V_1.5 anterograde trafficking was also investigated using a trafficking block and release assay^10^. Atrial myocytes overexpressing h-K_V_1.5-HA were treated with brefeldin A to prevent ER/Golgi transport and trafficking was restarted for 2 h before addition of sucrose (or control medium). As expected, quantification of surface HA staining (*i.e.* without permeabilization) relative to total HA (*i.e.* after permeabilization) showed that clathrin blockade results in reduced *de novo* delivery of K_V_1.5 channel (Figure S2) as clathrin is also involved in the active transport of cargo proteins at the Golgi exit en route to the plasma membrane^11^. Indeed, clathrin vesicles can be observed in the perinuclear region of freshly isolated cardiomyocytes (Figure S2B) and surrounding the Golgi apparatus as illustrated by electron microscopy images (Figure S2C). Then, the endogenous K_V_1.5-mediated current *I*_Kur_ was recorded in control and sucrose-treated myocytes by whole-cell patch-clamp. The potassium current density was significantly increased at several tested voltages in sucrose-treated myocytes (Figure 2D). Note that the cell capacitance was not significantly different between control and sucrose-treated cells (control: 42±5 pF, sucrose: 38±3 pF) showing that the surface membrane remains unchanged by the sucrose treatment. *I*_Kur_ was also recorded in myocytes in which the clathrin pathway was repressed using a small interfering RNA directed toward the clathrin heavy chain (si-clathrin HC). As observed with sucrose, the K_V_1.5-mediated current was significantly increased at several tested voltages in si-clathrin HC-transfected atrial myocytes (Figure 2E). Note that activation properties of *I*_Kur_ were not modified by clathrin blockade (Figure S3). Altogether, these results indicate that clathrin pathway is the major regulator of surface and functional expression of K_V_1.5 channels in atrial myocytes.

**Figure 2.**
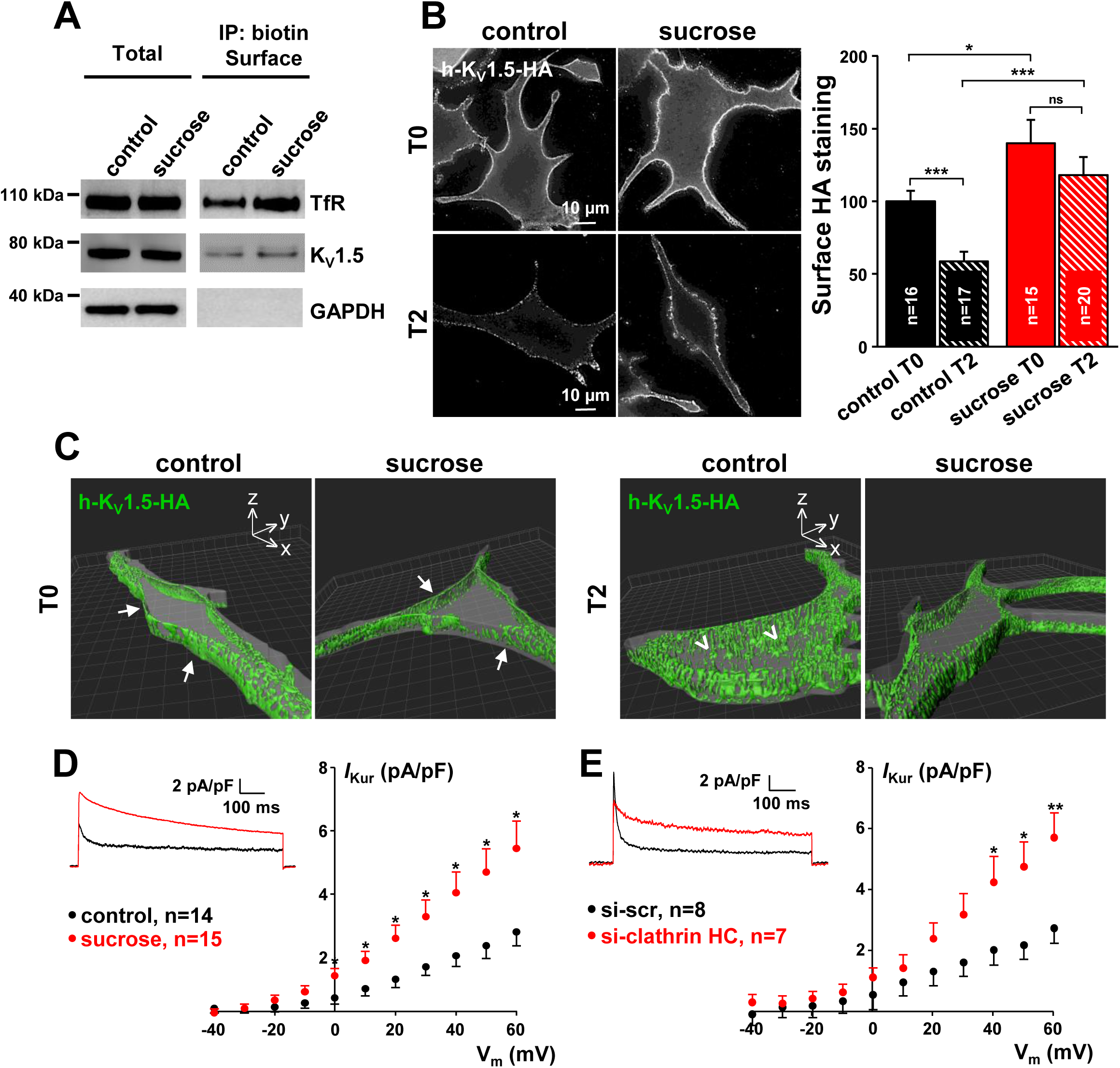
Blockade of clathrin pathway increases K_V_1.5 channel surface expression and corresponding *I*_Kur_ current in atrial myocytes. **A**, Representative surface biotinylation assays performed on atrial myocytes expressing h-K_V_1.5-GFP. Transferrin receptor (TfR) was used as a positive control of clathrin-dependent endocytosis. **B**, Surface expression and internalization of h-K_V_1.5-HA channel in control and sucrose conditions. (**Left**) Example images of cultured atrial myocytes live-stained with anti-HA and secondary Fab A488 antibody. Cells were immediately fixed (T0) or incubated for 2 hours before fixation (internalization, T2). (**Right**) Summary graphs of the fluorescent HA staining at cell surface at T0 and T2. **C**, Illustrative perspective views of 3D-rendered images (z-projections) at the two time points. Cell surface perimeters in grey have been done using Imaris’ surface module. Arrows point cell boundaries at T0; arrowheads point channels inside the cell at T2 in control condition. **D**, Current density-voltage relationships of endogenous *I*_kur_ recorded from atrial myocytes in control and sucrose conditions and representative traces in the insert (holding potential: −80 mV; test potential: +60 mV). **E**, Current density-voltage relationships of endogenous *I*_kur_ obtained from atrial myocytes transfected with si-scr or si-clathrin heavy-chain (HC) and representative traces in the insert. * P < 0.05; ** P < 0.01; *** P < 0.001, n=number of cells.

### Clathrin pathway inhibition leads to K_V_1.5 clusterization and association with stable microtubules

Movies of 2 hours were acquired from live atrial myocytes upon TIRF illumination to investigate the effect of clathrin blockade on K_V_1.5 channels dynamics in the membrane plane. In control condition, h-K_V_1.5-GFP channels showed a punctate distribution at the sarcolemma (vesicle mean size: 0.12±0.01µm^2^) (Figure 3A, E) with dynamic movements in the x*/*y axis (Movie 1A). Upon sucrose treatment, total evanescent field fluorescence (EFF) augmented linearly without reaching a plateau (Figure 3B) and was increased by ∼ 25% at the end of the movie (Figure 3C, P<0.001). Note that sucrose did not affect neither cell morphology nor GFP distribution used as control (Figures S4). Sucrose treatment also leaded to the accretion of h-K_V_1.5-GFP-containing vesicles into large clusters (cluster mean size: 2.77±0.05µm^2^) (Figure 3E, Movie 1B). The number of h-K_V_1.5-GFP vesicles having a diameter superior to 0.4 µm started to increase after 30 minutes of treatment and reached a plateau after 80 minutes (Figure 3D). These results suggest that the linear fluorescence increase is likely due to the continuous insertion of vesicles of smaller diameter from submembrane stores.

**Figure 3.**
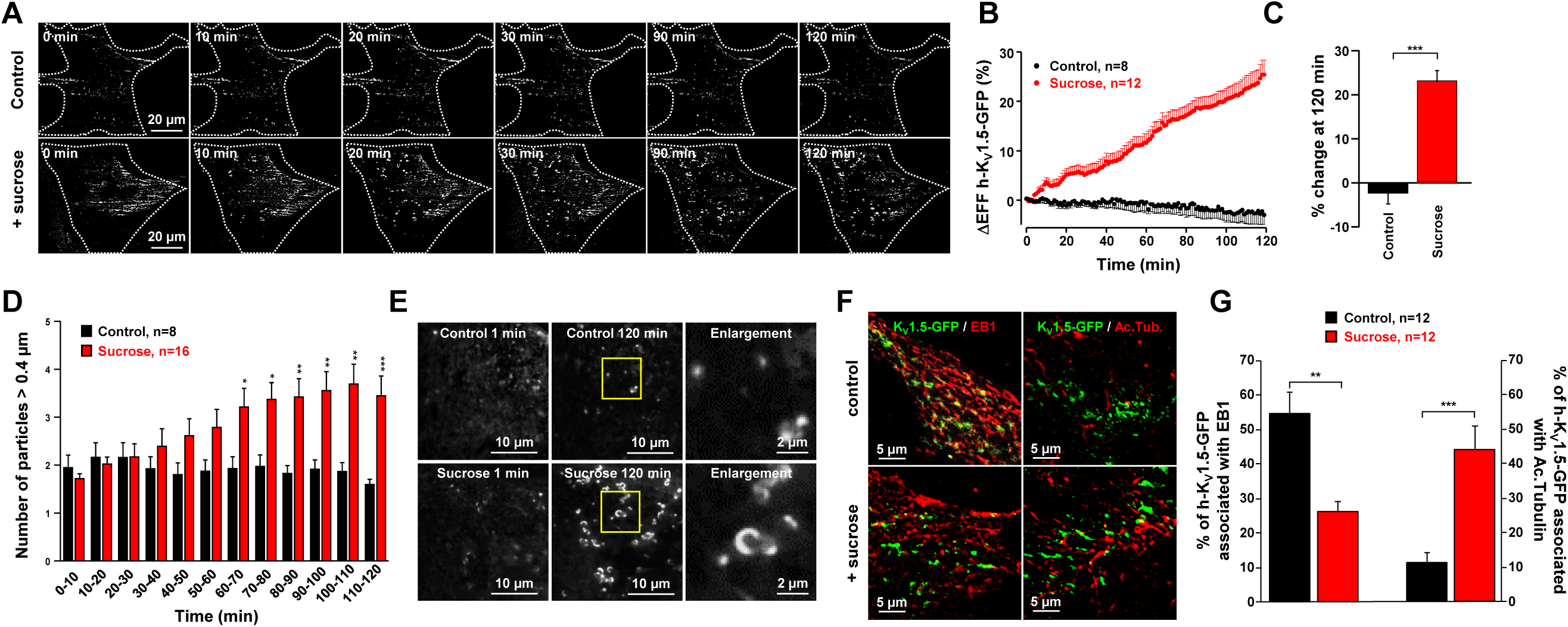
Blockade of the clathrin pathway induces K_V_1.5 clusterization in atrial myocytes. **A**, Example TIRF microscopy images from 2 hours time-lapse movie of cultured atrial myocytes expressing h-K_V_1.5-GFP under control (**top**) or sucrose (**bottom**) treatment. White dotted lines highlight cell boundaries delineated from corresponding DIC images. **B**, Mean time courses of total evanescent field fluorescence (EFF) intensity in response to sucrose treatment *vs* control. **C**, Bar graph of EFF intensity changes at the end of the movies. **D**, Bar graphs showing the number of h-K_V_1.5 channel vesicles over time in control and sucrose conditions. **E**, Example regions of interest (ROI) from TIRF movies at the beginning and at the end of the recording in control (**top**) and sucrose (**bottom**) conditions, and enlargements of outlined regions. **F**, Example TIRF images of control and sucrose-treated myocytes stained with EB1 (**left**) or acetylated tubulin (**right**). **G**, Bar graphs showing the association of h-K_V_1.5-GFP vesicles with EB1 or acetylated tubulin in control and sucrose conditions. * P<0.05; ** P<0.01; P<0.001; n=number of cells.

Large clusters observed in sucrose-treated cells were found to be associated with microtubule markers (Figure 3F). Whereas in control condition h-K_V_1.5-GFP vesicles were associated by 54.7±5.9% to dynamic microtubules stained with the plus-end tracking protein EB1, sucrose treatment reduced this association to 26.3±2.9% (Figure 3F, G). Conversely, large clusters were more associated with stables, acetylated microtubules (43.8±7%) compared to control h-K_V_1.5-GFP vesicles (11.4±2.7%) (Figure 3F, G). Thus, after clathrin blockade, h-K_V_1.5-GFP clusters are preferentially associated with stable microtubules.

### K_V_1.5 channel dynamics is reduced by clathrin blockade in atrial myocytes

Analysis of two hour movies acquired from live atrial myocytes in TIRF illumination also revealed that blockade of the clathrin pathway decreased K_V_1.5 channels mobility in the x/y axis as illustrated by Figure 4A, B (Movie 1C). To investigate further the effect of clathrin blockade on K_V_1.5 channels mobility in the membrane plane, dynamics of individual h-K_V_1.5-GFP vesicles was tracked from 2 minute movies in control and sucrose-treated myocytes. For the sake of clarity, single particles dynamics refer to track duration, track max speed, track displacement length, and track straightness (Figure 4C). In control condition, h-K_V_1.5-GFP channels displayed an average run of 3.89±0.34 µm in the x/y axis, and a mean duration of 24.5±3.1s. The max speed value was 1.05±0.04 µm/s (Table S1, Movie 1C). The behavior of h-K_V_1.5-GFP vesicles was also analyzed on the basis of the straightness of their tracks, 0 representing disordered runs and 1 representing 100% linear movement. In control condition the average track straightness was 0.65±0.03 (Table S1), suggesting a structured displacement in the membrane plane.

**Figure 4.**
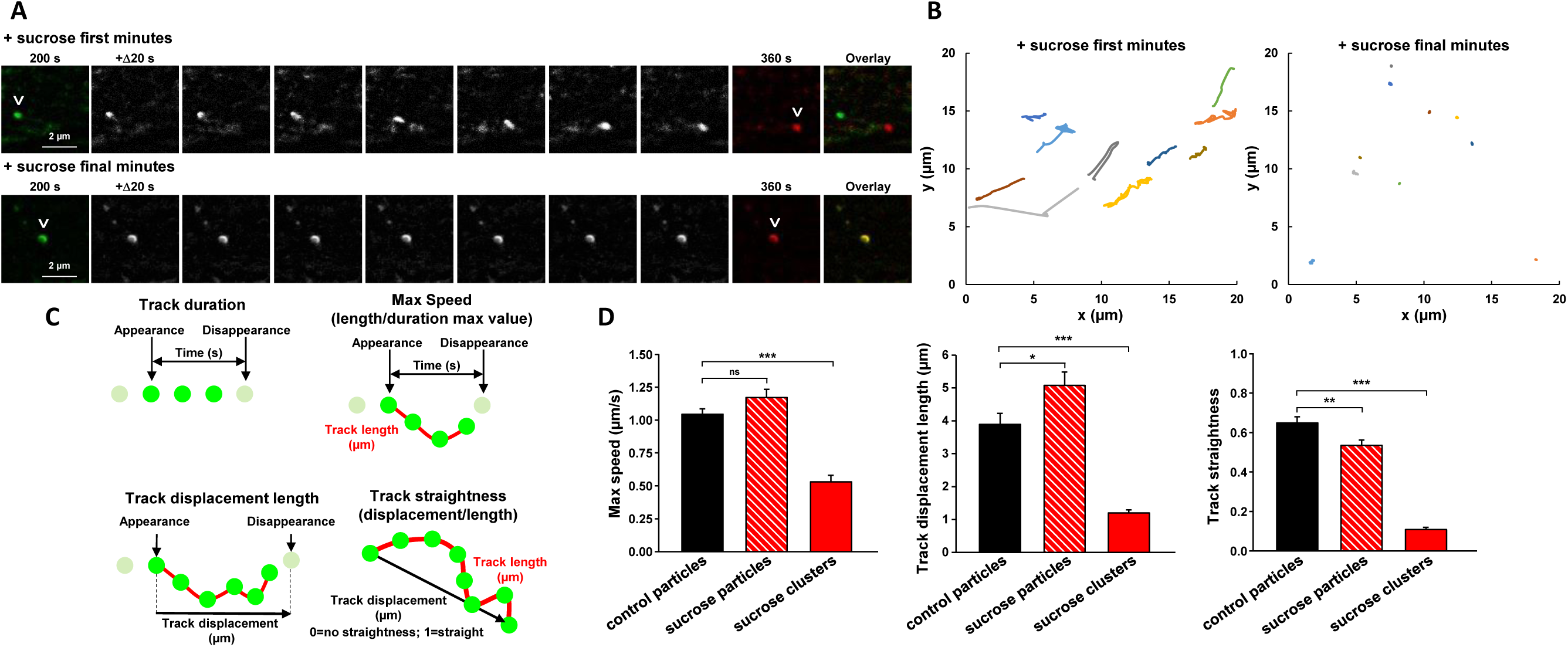
Blockade of the clathrin pathway decreases K_V_1.5 channels mobility in the sarcolemma of atrial myocytes. **A**, Example TIRF microscopy images of h-K_V_1.5-GFP vesicles displacements during first minutes (**top**) and final minutes (**bottom**) of sucrose treatment. Arrowheads point vesicles at the beginning and at the end of 160 s trackings. **B**, Example tracks generated from TIRF movies representing the displacement of h-K_V_1.5-GFP channels in the x/y axis during the first 10 minutes (**left**) and final 10 minutes (**right**) of a 2 h movie. **C**, Cartoons illustrating parameters measured from fast tracking movies. Green dots represent tracked h-K_V_1.5-vesicles (diameter ≥ 0.4 µm). **D**, Summary plots of h-K_V_1.5-EGFP channel dynamics in control and sucrose conditions. ns, not significant; * P < 0.05; *** P < 0.001; n=10-15 cells, 50-100 tracks/cell.

Clathrin blockade strongly affected the dynamics of h-K_V_1.5-GFP in the membrane plane, in particular in large sucrose-induced clusters that showed a profound mobility reduction. Large clusters persisted in the membrane plane for almost the whole duration of the recordings (100.6±3.3 s, Movie 1D), with an average displacement length of 1.19±0.1 µm and a track straightness of 0.11±0.01. Track max speed was also deeply affected, with an average value of 0.53±0.05 µm/s (Table S1). Please note that these values can be slightly underestimated (track duration) or overestimated (track max speed, and displacement length) as the tracking algorithm, which is designed to detect small fast moving particles, might have difficulties in following large slow moving clusters. Interestingly, sucrose-mediated clathrin blockade did not completely abolish the presence of moving particles. In fact, moving particles were tracked also after sucrose treatment and showed an increased persistence at the membrane plane compared to control: average displacement length 5.08±0.41 µm, average track duration of 38.6±3.4s (Table S1). Track straightness was also reduced (0.54±0.03) compared to control, while track max speed was not significantly affected (Table S1). Overall, this indicates that clathrin blockade reduced h-K_V_1.5-GFP mobility by inducing particles clusterization and increasing the particles/clusters persistence in the membrane plane.

### Role of the cytoskeleton in K_V_1.5 mobility in the sarcolemma

As h-K_V_1.5-GFP vesicles displayed a structured displacement in control atrial myocytes and were associated with stable microtubules after clathrin blockade, we investigated the association of the channel with the microtubule network and the actin cytoskeleton.

The effect of cytoskeleton disrupting agents was examined on K_V_1.5 total surface expression using whole-cell patch-clamp and live-cell immunostaining. Disruption of either microtubules with colchicine (2 h, 10 µmol/L) or actin cytoskeleton with cytochalasin D (24 h, 10 µmol/L) had no effect on endogenous *I*_kur_ in atrial myocytes (Figure S5A, B). Similarly, neither colchicine nor cytochalasin D significantly modified the total surface expression of the channel as quantified in atrial myocytes transduced with an extracellularly-HA-tagged h-K_V_1.5 channel^12^ (Figure S5C, D).

The effect of cytoskeleton disrupting agents on both anterograde and retrograde trafficking was then studied. To investigate *de novo* delivery of K_V_1.5 to the cell surface, we used the brefeldin A trafficking block and release assay developed by Smyth and colleagues^10^ (Figure S6A). Adult atrial myocytes transduced with h-K_V_1.5-HA were treated with BFA for 16 hours to block ER/Golgi export. After trafficking was restarted, disrupting agents were introduced for additional incubation time. HA surface staining was performed in live cell and total HA staining was performed after fixation and permeabilization. Images were acquired using high resolution 3-D deconvolution microscopy and the ratio between surface HA and total HA was quantified. Colchicine treatment reduced K_V_1.5 surface expression by 40.3±8.1% whereas cytochalasine D reduced it by 15.6±5.8%. To investigate the role of microtubules and actin cytoskeleton on K_V_1.5 retrograde trafficking, atrial myocytes transduced with h-K_V_1.5-HA were treated with either colchicine or cytochalasin D before been live-stained with an anti-HA antibody (Figure S6B, and S7). Some cells were immediately fixed (T=0) whereas others were proceeded for 2 hours internalization at 37°C (T=2h). Surface fluorescence intensity was quantified at cell boundaries using DIC images and normalized to control-treated cells. Colchicine and, to a lesser extent, cytochalasin D, reduced K_V_1.5 internalization compared to their respective control (surface staining EtOH: 30.8±9.6 *vs* colchicine: 49.5±8.4, −37%; surface staining DMSO: 42.3±9.3 *vs* cytochalasin D: 51.8±10.4, −18%).

Atrial myocytes transduced with h-K_V_1.5-GFP were labelled with the CellLight® Tubulin-RFP fluorescent dye and imaged by TIRF microscopy using dual color acquisitions in real time. In the sarcolemma plane, h-K_V_1.5-GFP vesicles followed the microtubule network as shown in Figure 5A. The mobility pattern of h-K_V_1.5-GFP vesicles was analyzed using the standard deviation (SD) projection from a time series. The resulting SD map was aligned with the average (fixed) tubulin image. K_V_1.5 vesicles exhibited a linear movement that followed the microtubule network (Figure 5B). TIRF movies were acquired for 2 hours from atrial myocytes treated with the microtubule disrupting agent colchicine (10 µmol/L) (Movie 2A). The h-K_V_1.5-GFP EFF signal was stable for 40 minutes, then the number of h-K_V_1.5-GFP vesicles (> 0.4 µm) increased by 39% in the sarcolemma after 40 minutes and reached a maximum increase of 74% after 1-hour colchicine treatment (Figure 5C). After 2 hours treatment with colchicine, the particles mobility was severely compromised in the x/y axis (Figure 5D). In addition, 2 minutes TIRF movies were acquired in order to measure single particles dynamics in control (EtOH) and in the presence of colchicine (Movie 2B). In fact, over a total of 10 acquired movies only 4 moving particles were detected, while 43 moving particles were tracked over 9 movies in control. In addition, the mobility of those few moving particles was strongly affected by the presence of colchicine. In particular, track duration was increased to 24.0±7.1 s (*vs* 13.4±1.6 s in control) and track displacement length was reduced to 0.99±0.21 µm (*vs* 4.17±0.37 µm in control). Track straightness and track max speed were also all significantly decreased compare to control (Figure 5E, Table S1).

**Figure 5.**
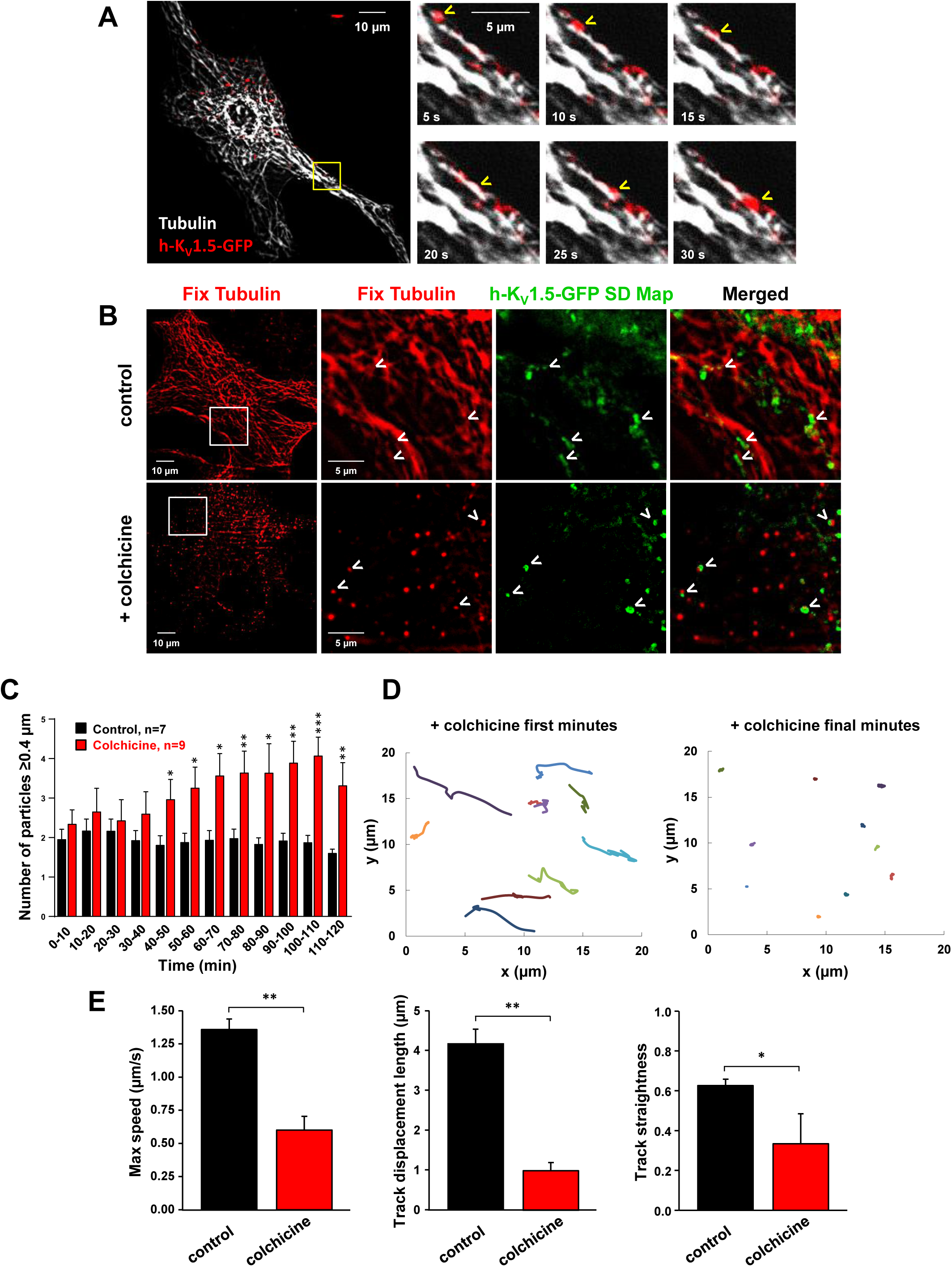
Microtubules disruption reduces K_V_1.5 channel mobility in the sarcolemma of atrial myocytes. **A**, Example TIRF microscopy images of dual color acquisitions from a time series movie showing h-K_V_1.5-GFP particles (red) and microtubules (white), and enlargement of one ROI. **B**, Standard deviation (SD z-projection) maps of h-K_V_1.5-GFP vesicle tracks compared to fixed cell images (average z-projection) of microtubules. Arrowheads indicate displacement (or not) of h-K_V_1.5-GFP vesicles (SD maps) in control and colchicine-treated myocytes. **C**, Bar graphs showing the number of h-K_V_1.5-GFP channel particles over time in control and colchicine conditions. **D**, Example tracks generated from TIRF movie illustrating the displacement of h-K_V_1.5-GFP channels in the x/y axis during first (**left**) and final (**right**) minutes of colchicine treatment. **E**, Summary plots of h-K_V_1.5-EGFP channel dynamics from fast tracking TIRF movies in control and colchicine conditions. * P < 0.05; ** P < 0.01; *** P < 0.001; n=5-15 cells, 50-100 tracks/cell.

Similar experiments were repeated in atrial myocytes transduced with h-K_V_1.5-GFP to investigate the role of the actin cytoskeleton. Cells were labelled with the CellLight® Actin-RFP fluorescent dye and imaged in dual color acquisitions by TIRF microscopy. As shown in Figure 6A, h-K_V_1.5-GFP vesicles did not track along the actin cytoskeleton in the membrane plane but rather moved between actin fibers. We examined the mobility pattern of h-K_V_1.5-GFP vesicles in relation with the actin cytoskeleton. The standard deviation projection of h-K_V_1.5-GFP vesicles from a time series was aligned with the average (fixed) actin image. Linear movements of h-K_V_1.5-GFP particles close to the actin cytoskeleton but not overlapping it were observed (Figure 6B). After overnight treatment with an inhibitor of actin polymerization, cytochalasin D (10 µmol/L), the particles mobility was not massively compromised in the x/y axis (Figure 6C, Movie 3A). To test the involvement of the actin cytoskeleton in h-K_V_1.5-GFP vesicles dynamics in the membrane plane, 2 minutes long TIRF movies were acquired from atrial myocytes (Movie 3B). The treatment with cytochalasin D did not significantly affect h-K_V_1.5-GFP particles dynamics. In particular, the average track displacement length was 4.00±0.41 µm in control and 3.24±0.38 µm in the presence of cytochalasin D, with an average track duration of 13.86±1.49 s and 11.61±1.57 s, respectively (Figure 6D, Table S1). Track max speed and track straightness were also unaffected by the treatment with cytochalasin D. However, the total number of moving particles was reduced by the treatment. In fact, a total of 43 particles were tracked over 10 movies in control condition, while, in the presence of cytochalasin D, it was possible to track only 18 moving particles over 9 acquired movies. This might indicate a partial involvement of the actin cytoskeleton in the membrane plane.

**Figure 6.**
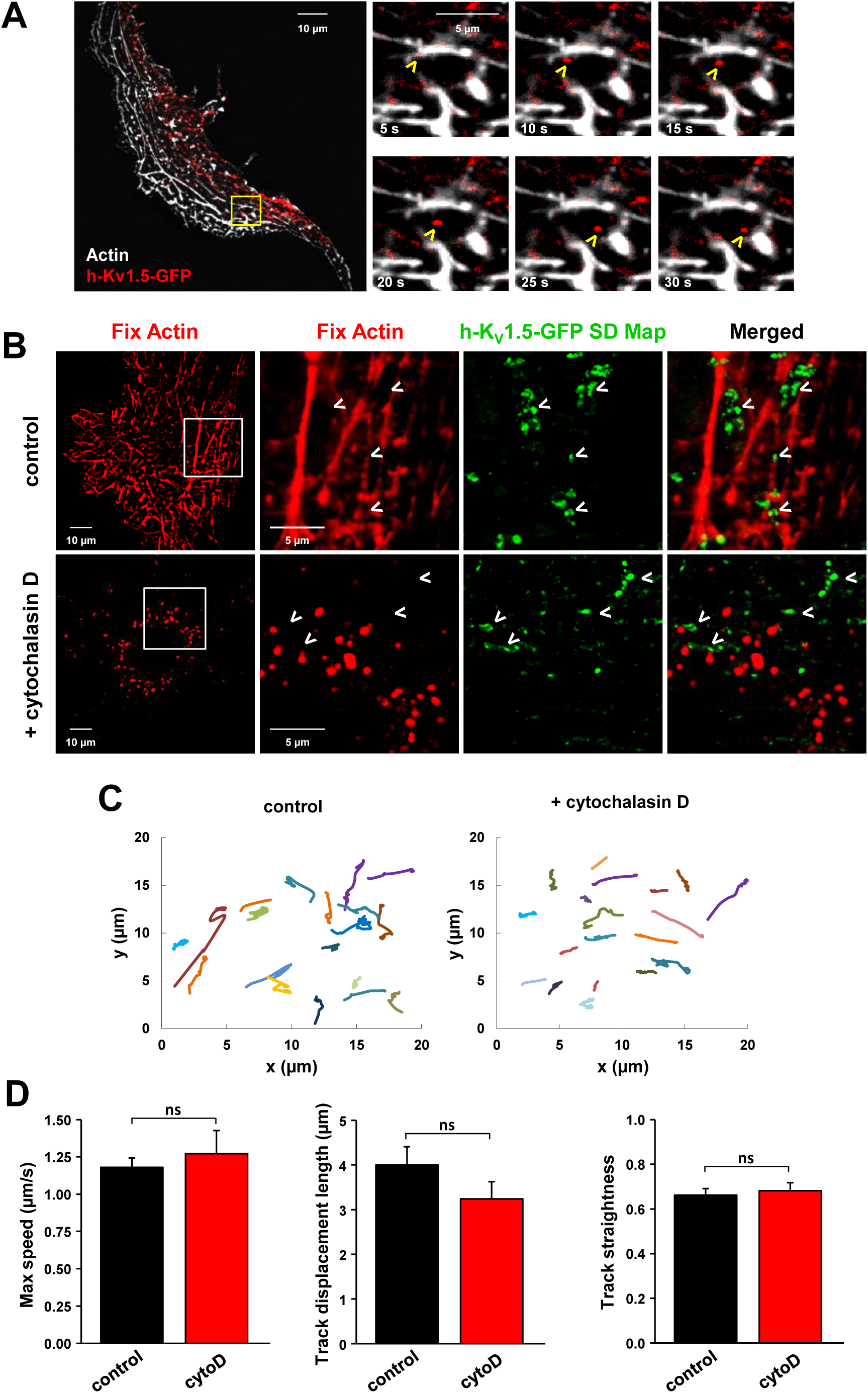
Actin disruption does not affect K_V_1.5 channel mobility in the sarcolemma of atrial myocytes. **A**, Example TIRF microscopy images of dual color acquisitions from a time series movie showing h-K_V_1.5-GFP particles (red) and actin (white), and enlargement of one ROI. **B**, Standard deviation (SD z-projection) maps of h-K_V_1.5-GFP vesicle tracks compared to fixed cell images (average z-projection) of microtubules. Arrowheads indicate displacement of h-K_V_1.5-GFP vesicles (SD maps) in control and cytochalasin D-treated myocytes. **C**, Example tracks generated from 10 minutes time-lapse TIRF movies illustrating the displacement of h-K_V_1.5-GFP channels in the x/y axis in control (**left**) or and cytochalasin D-treated myocytes (**right**). **D**, Summary plots of h-K_V_1.5-EGFP channel dynamics from fast tracking TIRF movies in control and cytochalasin D conditions. ns, not significant; n=20-40 cells, 50-100 tracks/cell.

Quantifications of h-K_V_1.5-GFP vesicles associated with tubulin or with actin in the membrane plane showed that K_V_1.5 was twofold more associated with microtubules, than with actin fibers in control conditions (∼36% *vs* ∼18%) (Figure 7A, B). After colchicine treatment, the percentage of h-K_V_1.5-GFP associated with microtubule remnants was reduced by half whereas the association h-K_V_1.5-GFP with actin residues was not significantly modified by the cytochalasin D treatment (Figure 7B). Altogether, these results support the preferential involvement of microtubules over actin in K_V_1.5 dynamics in the membrane plane in atrial myocytes. Finally, in order to identify the large clusters observed after clathrin blockade, several antibodies were tested and only tubulin markers were detected as positives. Interestingly, whereas in control conditions h-K_V_1.5-GFP vesicles were associated by 54.7±5.9% to dynamic microtubules stained with the plus-end tracking protein EB1, sucrose treatment reduced this association to 26.3±2.9% (Figure 7C, D). Conversely, h-K_V_1.5-GFP were more associated with acetylated (i.e. stables) microtubules when treated with sucrose (43.8±7%) compared to control (11.4±2.7%) (Figure 7C, D). Thus, after clathrin blockade, h-K_V_1.5-GFP clusters are preferentially associated with stable microtubules.

**Figure 7.**
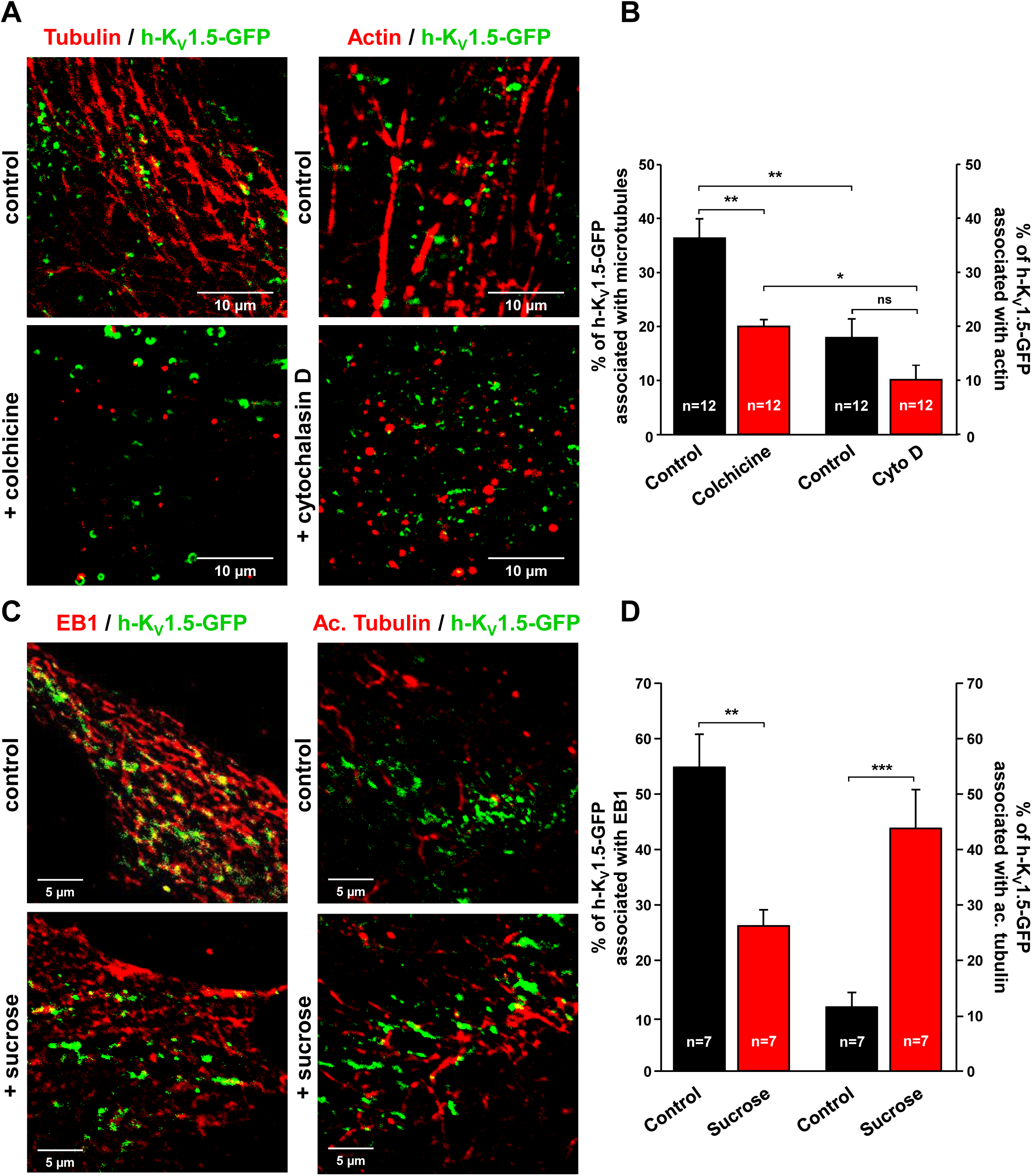
Association of h-K_V_1.5-GFP with the cytoskeleton. **A**, Example ROIs from microscopy images of cultured atrial myocytes expressing h-K_V_1.5-GFP and stained with tubulin or actin in control or after treatment with cytoskeleton-disrupting agents colchicine or cytochalasin D. **B**, Summary plots showing the association between h-K_V_1.5-GFP channels and cytoskeleton markers in control and cytoskeleton-disrupting agents-treated cells. The number of green fluorescent vesicles positives for actin or microtubule markers were quantified and reported to the total number of green fluorescent vesicles. **C**, Example ROIs from microscopy images of cultured atrial myocytes expressing h-K_V_1.5-GFP and stained with dynamic (EB1-positives) or stable (acetylated tubulin-positives) microtubules in control and sucrose conditions. **D**, Summary plots showing the association of h-K_V_1.5-GFP vesicles with EB1 or acetylated tubulin in control and sucrose conditions. ns, not significant; * P < 0.05; ** P < 0.01; *** P<0.01; n=number of cells.

### Alteration of trafficking pathways in atrial fibrillation

Finally, we examined K_V_1.5 channel trafficking pathways during AF. First, several trafficking-related proteins were quantified using western blot assays performed with total lysate extracted from permanent AF (n=4-5) and control (sinus rhythm, SR, n=5) patients (Figure 8A, B). K_V_1.5 total protein expression level was increased by 35.2±7.1% in permanent AF patients compared to SR controls. A marked increase in effectors related to trafficking was observed in AF such as endocytosis markers (clathrin heavy-chain, CHC: +49.1±14.8% and dynamin-2: +111.8±13.3%), retrograde molecular motors (myosin 6: +89.0±12.8% and dynein: +31.2±9.4%), proteins associated with early endosomes (EEA1: +69.3±13.9% and Rab5: +53.6±8.4%), and finally recycling-endosome associated proteins (Rab-4: +103.4±9.1% and Rab11: +64.3±6.1%). Moreover, while the total expression level of tubulin was not modified in permanent AF, the expression of dynamic microtubules was increased (plus-end tracking protein EB1: +37±5%) whereas stable microtubules expression was decreased (acetylated tubulin: −40.0±8.2%) compared to control.

**Figure 8.**
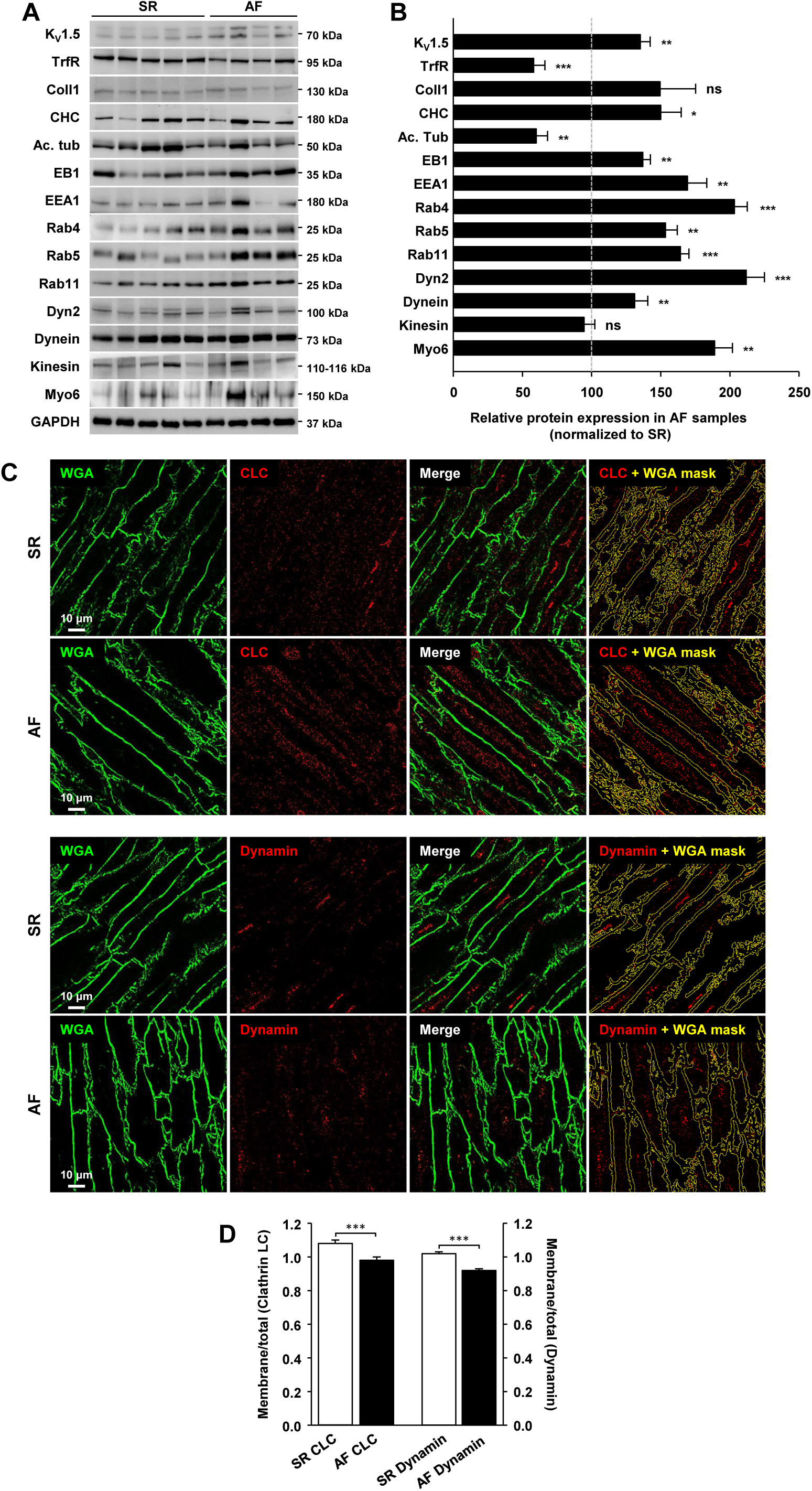
Trafficking-related proteins are altered during atrial fibrillation. **A**, Representative western blots showing expression levels of K_V_1.5, transferrin receptor, collagen 1 and trafficking-related markers in whole atrial lysates from control (sinus rhythm, SR) and atrial fibrillation patients (AF). GAPDH served as a loading control. **B**, Corresponding histograms showing expression levels of proteins normalized to GAPDH and expressed in % changes from SR samples. N=3-10 experimental replicates; n=4-5 patients/group. **C**, Representative microscopy images showing the distribution of clathrin light-chain (CLC) and dynamin in SR and AF tissue samples. The membrane marker WGA (left) was used to generate a mask (right) delineating myocytes membranes. **D**, Summary bar graph showing the distribution of CLC and dynamin signals in the membrane over total fluorescence distribution. N=24-25 images/patient; n=3 patients/group. ns, not significant; * P<0.05; ** P<0.01; *** P<0.001.

We then analyzed the distribution of two selected endocytosis markers in the atria of AF patients. Frozen slices from biopsy samples of three AF patients and three SR controls were stained with antibodies against clathrin light-chain (CLC) and dynamin-2. Wheat Germ Agglutinin (WGA) was used in order to stain cardiomyocyte membranes. Series of 8-9 images for each sample were then acquired using high resolution 3-D deconvolution microscopy. The acquired images were then analyzed in Fiji where an algorithm was designed in order to calculate the ratio between “membrane mean intensity” and “total mean intensity” for each selected marker (Figure 8C, D). Both markers were almost evenly distributed within the control tissue, as the signal mean intensity was just slightly higher in the membrane region (membrane/total ratio 1.08±0.02 and 1.02±0.01 for CLC and dynamin-2 respectively). Interestingly, in AF condition a small but significant ∼10% decrease in the membrane/total ratio was observed for both CLC and dynamin-2 (0.98±0.02 and 0.92±0.01 respectively).

Altogether, these results indicate that the trafficking machinery is profoundly altered during permanent AF.

## DISCUSSION

The present study reveals the extreme mobility of K_V_1.5 channels in atrial cardiomyocytes that oscillate between a mobile/trafficking state and a fixed one that determine the level of protein expression at the sarcolemma. The mobility of K_V_1.5 channels depends on the integrity of clathrin-mediated endocytosis pathway and microtubule network. During AF, both retrograde and anterograde trafficking pathways are altered pointing to a possible shift towards enhanced mobile state of K_V_1.5 channels in the sarcolemma of diseased atrial myocytes.

### Clathrin-mediated endocytosis of KV1.5 channel in native atrial myocytes

Previous studies have reported dynamin-dependent internalization of K_V_1.5, but the precise role of the clathrin- or caveolin-dependent pathways were not elucidated. Whereas K_V_1.5 has been shown to target caveolae in Ltk2 and FRT cells stably expressing K_V_1.5^7,8^, a very limited association between K_V_1.5 and caveolin 3 has been observed in rat and canine hearts^13^. Using a step sucrose gradient, we previously showed that K_V_1.5 and caveolin 3 are found in distinct fractions of proteins extracted from rat atrial myocardium^14^. Our present findings confirm that K_V_1.5 channels are not associated with caveolin 3 in atrial myocytes but preferentially associated to clathrin vesicles.

Preventing clathrin endocytosis using sucrose treatment triggered the accumulation of K_V_1.5 channels in the sarcolemma and was associated with a marked enhancement of *I*_Kur_. Sucrose is known to induce the formation of clathrin microcages^9^, resulting in the observed clusterization and immobilization of K_V_1.5 in the membrane plane. The immobilization and clustering of K_V_1.5 in atrial myocytes have been already described and was attributed to the anchoring of channels by the MAGUK protein SAP97 in microdomains of the sarcolemma^15^. A similar immobilization of K_V_ channels in membrane subdomains has been reported for neuronal K_V_2.1 subunits. K_V_2.1 channel forms large clusters on the plasma membrane of hippocampal neurons and transfected HEK cells where they remain immobile after delivery at the insertion site^16^. Interestingly, clustered K_V_2.1 channels do not efficiently conduct K^+^, whereas the non-clustered channels are responsible for the K_V_2.1-mediated current^17^. In addition, we observed that sucrose-mediated clathrin inhibition also increased the track straightness and track displacement length of the non-clustered K_V_1.5 vesicles moving in the membrane proximity. This suggests an increased persistence of K_V_1.5 vesicles in the membrane plane, consistent with a reduction of channel internalization. In the present study, the observation of an increase in *I*_Kur_together with an increase in K_V_1.5 cell surface expression, clusterization and vesicles persistence in the membrane plane showed that K_V_1.5 channels are functional after clathrin blockade.

### Preferential association of K_V_1.5 channel with the microtubule network

Both the microtubule network and actin filaments are involved in ion channel retrograde and anterograde trafficking and cytoskeleton damages have been shown to impact ion channel expression.

For instance, alteration of microtubule integrity with microtubule depolymerizing agents has a profound effect on the surface expression of both potassium and sodium channels. In atrial myocytes, nocodazole treatment increases the surface expression of K_V_1.5 channels and the amplitude of the corresponding current, *I*_Kur_^5^. Microtubule disruption by colcemid treatment was shown to impair the normal surface distribution of K_V_1.5 by increasing its internalization in L-cells^7^. Similar results were reported for K_V_4.2 and hERG channels, the molecular basis of *I*_to_ and *I*_Kr_ respectively. Treatment with the dynein inhibitor erythro-9-(2-hydroxy-3-nonyl)adenine hydrochloride or overexpression of the p50 protein, which leads to the dissociation of the dynactin/dynein complex, reproduced the effect of microtubule disruption by nocodazole in HEK cells^18^. In neonatal rat cardiomyocytes, stabilization of the microtubule cytoskeleton reduces *I*_Na_ density via a mechanism that reduces channel expression at the sarcolemma^19^. In HL-1 cells, anterograde trafficking of K_V_1.5 channel has been shown to occur preferentially on dynamic microtubules associated with the kinesin-2 motor, KIF17^20^. Therefore, microtubules are important determinants of ion channel trafficking in both the retrograde and anterograde direction.

Microtubules can be classified on the basis of their dynamic properties, distinguishing stable *vs* dynamic microtubules. With the exception of a small stable population, microtubules are in a state of dynamic instability with addition of subunits to the plus end and loss at the minus end^21^. In the present study, microtubule depolymerization mimicked the effect of the clathrin blockade. Indeed, colchicine induced the accumulation and the immobilization of K_V_1.5 channels in the membrane plane. The protein EB1 interacts indirectly with the dynein/dynactin complex, the microtubule retrograde motor^22^. Whereas K_V_1.5 channels were strongly associated with EB1-positives dynamic microtubules in control conditions, K_V_1.5 channels were preferentially associated with acetylated microtubules after clathrin blockade.

Taken together, these results support a major role for microtubules in both the internalization and the mobility of K_V_1.5 channels in native atrial myocytes.

### Moderate involvement of the actin cytoskeleton in K_V_1.5 channel dynamics and internalization

In hippocampal neurons, treatment with latrunculin-A, which prevent actin polymerization, induces a redistribution of K_V_2.1 channels from the cell body to the entire cell, together with an increase in cluster size and a decrease in cluster number, suggesting a role for cortical actin in both cluster maintenance and localization of K^+^ channels in restricted areas^16^. Manipulations of the actin cytoskeleton also affects ion channel function by affecting channel trafficking. For instance, K_V_1.5 surface localization has been shown to depend on an intact actin cytoskeleton as indicated by the increase of K_V_1.5 at the surface of HL-1 cardiac cell line treated with cytochalasin D^24^. Both actin depolymerization and antisense RNA directed against α-actinin2 increase the K_V_1.5-mediated current and the number of active channels in HEK293 cells. From these observations, it has been suggested that actin cytoskeleton disruption favors the release of intracellular pools of channels from anchors and channel insertion into the plasma membrane^25^. Similarly, whereas the delivery of connexin-43 at the intercalated disc was shown to be directly mediated by microtubules, actin-associated connexin-43 functions as an alternative trafficking route, providing a cytoplasmic reservoir recruitable in response to cellular stress ^10,22^.

In the present study, disruption of the actin cytoskeleton did not change the surface expression of K_V_1.5 channel in the membrane plane. Also, K_V_1.5 particles dynamics (displacement length, track duration and straightness, and max speed) were not affected by the treatment with cytochalasin D. Only the total number of detectable moving particles in the membrane plane was decreased. Since an indirect linkage to the actin cytoskeleton via α-actinin2 have been reported for K_V_1.5^25^, one could hypothesize that the decrease in detectable moving particles after cytochalasin D treatment affect a binding protein that participates in labile/transitory maintenance of K_V_1.5 channel in the membrane plane. At the moment, the role of actin binding proteins on K_V_1.5 channel trafficking in the membrane plane in atrial myocytes sarcolemma remains to be investigated.

Taken together these results indicate that the actin cytoskeleton has a limited role in K_V_1.5 channel trafficking in native atrial myocytes. This conclusion is also supported by our previous observation showing that K_V_1.5 channel delivery to the sarcolemma was more dependent on the microtubule network than on cortical actin cytoskeleton^4^.

### Physiological relevance during permanent atrial fibrillation

Changes in channel expression are now recognized as the main mechanism underlying the altered electrical properties of diseased myocardium. Furthermore, during AF, K_V_1.5 channel expression varies with arrhythmia duration, as indicated by the observation of a positive correlation between channel protein density and the duration of the sinus rhythm after the last paroxysmal AF episode^26^. Similarly, a fast and transient increase in K_V_1.5 channel expression has been observed during paroxysmal atrial tachycardia in rats^27^.

The present study suggests that K_V_1.5 channel dynamics also contribute to the complex pattern of K_V_1.5 channel expression during disease progression. Our finding of up-regulated recycling pathways and reduced clathrin-mediated endocytosis is consistent with an accumulation of K_V_1.5 channels in the sarcolemma of atrial myocytes from patients in permanent AF. We previously reported that chronic hemodynamic overload of the atria, an AF risk factor, stimulates K_V_1.5 channel recycling and exocytosis^4^. Finally, alteration of microtubule network states as observed in remodeled atrial myocytes could also impact K_V_1.5 channel properties.

Thus, the machinery identified in this study provides mechanistic insight into how K_V_1.5 dynamics could play major role in adaption of atrial electrical properties to modifications of mechanical and/or homeostatic states of the atrial myocardium, whether in physiological or pathological settings^3,4,11^.

## MATERIALS AND METHODS

### Animals and cardiac myocytes isolation

Adult male Wistar rats (Janvier) were treated in accordance with French Institutional guidelines (Ministère de l’Agriculture, France, authorization 75-1090) and conformed to the Directive 2010/63/EU of the European Parliament. Adult atrial myocytes were obtained by enzymatic dissociation on a Langendorff column as previously described^4^. Briefly, hearts were cannulated and retrogradely perfused at 5mL/min through the aorta, first with a solution containing (in mmol/L): NaCl 100, KCl 4, MgCl_2_ 1.7, glucose 10, NaHCO_3_ 5.5, KH_2_PO_4_ 1, HEPES 22, BDM 15, taurine 10, pH 7.4 (NaOH) at 37°C for 5min, then with an enzymatic solution containing (in mmol/L): NaCl 100, KCl 4, MgCl_2_ 1.7, glucose 10, NaHCO_3_ 5.5, KH_2_PO_4_ 1, HEPES 22, BDM 15 and 200UI/mL collagenase A (Roche Diagnostics) plus 0.5% bovine serum albumin, pH 7.4 (NaOH) at 37°C for ∼17min. Solutions were bubbled with 95% O_2_/5% CO_2_ throughout. Atria were separated and placed into Kraft-Brühe (KB) buffer containing (in mmol/L): glutamic acid potassium salt monohydrate 70, KCl 25, taurine 20, KH_2_PO_4_ 20, MgCl_2_ 3, EGTA 0.5, glucose 10, HEPES 10, pH 7.4 (KOH). The atria was then cut into small pieces and gently agitated to dissociate single myocytes. Myocytes were plated onto laminin-coated petri dishes and cultured in standard conditions (37°C, 5% CO_2_) in M199 medium containing 5% FCS, 1‰ ITS and 1% P/S.

### Adenoviruses and cytoskeleton tagging

Human K_V_1.5 channel cDNA with the EGFP sequence inserted downstream the S6 segment sequence or with the HA tag into the coding sequence of the extracellular loop between residues 307 and 308 were subcloned in adenovirus. Rat adult atrial myocytes were transduced 24h after isolation with 3.7^10^6^ particles/mL of Ad-h-K_V_1.5-EGFP or with 3.7^10^6^ particles/mL of Ad-h-K_V_1.5-HA. pcDNA3-K_V_1.5-HA (Hemagglutinin) was generated by inserting the coding sequence for the nine-amino-acid residue HA tag flanked by single glycine codons into the coding sequence of K_V_1.5 between extracellular residues 307 and 308 (Gift from D. Fedida). SiRNA sequence for clathrin heavy-chain Cy3 5’-uucuacaaacuagcaguaagg-3’ (si-CHC) or its control scramble (siscr) Cy3 5’-aauucuccgaacgugucacgutt-3’ were provided by MWG. Actin filaments and microtubules were stained with RFP fluorescent dyes using BacMam technology (CellLight®, Thermofisher, 7.5^10^5^ particles/mL). All studies were performed between 3-4 days post-transduction or transfection.

### Electrophysiological measurements

*I*_Kur_ properties were studied using standard whole-cell patch clamp. Pipette solution contained in mmol/L: K-aspartate 115, KCl 10, KH_2_PO_4_ 2, MgCl_2_ 3, HEPES 10, glucose 10, CaCl_2_ 0.1 and Mg-ATP 5, pH 7.2 (KOH). Cells were perfused with a solution containing in mmol/L: NaCl 140, KCl 4, MgCl_2_ 2, NaHPO_4_ 1, glucose 20, HEPES 10, CaCl_2_ 2, pH 7.4 (NaOH). For the current-V_m_ protocol, currents were elicited by 750ms depolarizing pulses from a holding potential of −80 mV, in 10 mV increments, between −130 and +60 mV, at a frequency of 0.2 Hz. Leak current was numerically compensated.

### Surface biotinylation assays

Live cells were surface biotinylated for 30 minutes at 4°C using PBS containing 0.5 mg/ml of EZ Link Sulfo-NHS-SS-Biotin (Sigma). Cells were washed with PBS and incubated for 10 minutes in PBS containing 100 mmol/L glycine to quench unbound biotin. Cells were then lysed for 30 minutes at 4°C with RIPA buffer containing, in mmol/L: NaCl 150 and Tris-HCl 50 (pH7.5) supplemented with 1% NP-40, 0.5% Na-deoxycholate, 0.1% SDS, 1% Triton X-100 and protease inhibitor cocktail (Sigma). After centrifugation, supernatants were incubated with 50 µl of streptavidin-sepharose beads (Thermofisher Scientific) overnight at 4°C. After five washes with RIPA buffer, samples were warmed for 20 minutes at 37°C in 2X sample buffer and analyzed by western blot.

### Western Blot

Proteins were separated on NuPAGE® Novex® 10% Bis-Tris gels (Life Technologies) and transferred on nitrocellulose membrane (Biorad). Incubation were performed with appropriate primary antibodies followed by HRP-coupled secondary antibodies (Cell Signaling). Proteins were revealed with ECL Prime (GE Healthcare) and images were acquired using a LAS4000 Camera (GE Healthcare).

### Immunofluorescence and high-resolution 3-D deconvolution microscopy

Immunofluorescence was performed on cardiomyocytes or on atrial cryosections (7μm-thick) fixed with 4% paraformaldehyde (PFA) for 10 min at room temperature (RT). Preparations were incubated for 1h at RT with permeabilizing/blocking buffer (PBS containing 0.1% Triton X-100, 1% BSA, 10% sera) and then incubated with primary antibodies diluted in blocking buffer (PBS containing 1% BSA, 3% sera) overnight at 4°C. Detection was performed with a 1h incubation at RT with secondary antibodies AlexaFluor^488^ or AlexaFluor^594^ (1:500; Molecular Probes), and DAPI (1:500; Sigma) to detect nuclei. Epifluorescence imaging was performed using a Plan APO 60X Oil, 1.42 NA objective in an Olympus IX-71 microscope. Digital images were captured with a sCMOS camera (Hamamatsu) in the Z-axis in 0.2 µm increments using advanced Nanomover technology. Images were deconvoluted using acquired PSF with DeltaVison Elite microscopy system (GE Healthcare).

### Anterograde trafficking and internalization assays

Brefeldin A trafficking block and release assay developed by Smyth and colleagues^10^ was used to follow d*e novo* delivery of K_V_1.5 channels to the cell surface. Adult atrial myocytes transduced with h-K_V_1.5-HA were treated with 2.5 µg/mL brefeldin A for 16 hours to block ER/Golgi export. Brefeldin A was removed with warm PBS, medium replaced and cells were incubated for additional 2 hours to restart trafficking. Cytoskeleton disrupting agents or control vehicles were introduced for additional incubation time: 2 hours for control EtOH and colchicine (10 µmol/L), 24 hours for control DMSO and cytochalasin D (10 µmol/L). HA-tag surface staining was performed on ice on live cells using anti-HA antibody. After staining with a AlexaFluor 488, cells were fixed and permeabilized to label total HA using anti-HA antibody and AlexaFluor 594. Images were acquired using high resolution 3-D deconvolution microscopy and the ratio between surface HA and total HA was quantified using ImageJ.

For internalization, myocytes transduced with h-K_V_1.5-HA were treated for 16 hours with 2.5 µg/mL brefeldin A to block ER/Golgi export prior to being treated with cytoskeleton disrupting agents or control vehicles in presence of brefeldin A: 2 hours for control EtOH and colchicine (10 µmol/L), 24 hours for DMSO and cytochalasin D (10 µmol/L). Cells were live stained on ice using anti-HA antibody and Fab fragment AlexaFluor 488. Cells were either immediately fixed and imaged (T0) or returned back to the incubator for additional 2 hours (T2, internalization). Images were acquired using high resolution 3-D deconvolution microscopy and surface fluorescence intensity was quantified at cell boundaries using corresponding DIC images and normalized to control-treated cells.

### Evanescent field microscopy and image analysis

Live imaging experiments were performed at ∼27°C with cultured myocytes seeded on glass bottomed micro-dishes (170µm thickness, Ibidi) bathed with standard extracellular solution. Kv1.5-EGFP channel dynamics was visualized with total internal reflection fluorescence (TIRF) microscopy using the Olympus Cell^tirf^ system. Cells were placed on an Olympus IX81-ZDC2 inverted microscope and TIRF illumination was achieved with the motorized integrated TIRF illuminator combiner (MITICO-2L) and dual band optical filter (M2TIR488-561) using a 60X/1.49 APON OTIRF objective, allowing GFP and RFP visualization. All image acquisitions, TIRF angle adjustment and some of the analysis were performed using the xcellence software (Olympus). Time series images were recorded using a digital CCD camera ORCA-ER (Hamamatsu). Cells were imaged for 10min before applying drugs to establish the baseline whole-cell evanescent field fluorescence (EFF). To quantify changes in EFF, the cell perimeter was delineated and the projection of minimum intensity (EFF_min_) of the whole movie was subtracted from each image (EFF_(t)_) and normalized to the background intensity measured in a region of interest outside the cell (EFF_bck_) using the following equation:

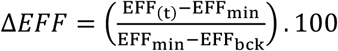

### Vesicle tracking and Dynamics parameters

In a first set of experiments, single particle tracks were created manually from consecutive frames in 20×20 µm regions of interest (ROI) from TIRF movies using the tracking Brownian motion model, which finds the most likely path connection in any direction. The minimum particle size diameter was fixed to 0.4 µm and the measured parameters included number of particles in the ROI, particles accumulation in the membrane plane over time and particles coordinates (in the x/y axis). Time series of 360 images at 20s intervals were acquired for control, sucrose and colchicine movies; time series of 150 images at 5s intervals were acquired for control and cytochalasin D movies. In addition, a second set of experiments was performed in order to obtain accurate measurements of single particles dynamics. In this case, time series of 240 images every 0.5s (total movie length of 2 min) were acquired for all the conditions listed above. Particles that exhibited a clear linear movement in the membrane plane were analyzed using the autoregressive motion algorithm of Imaris 9 and the following parameters were quantified: track duration, track displacement length, maximal speed (speed max in the text), and track straightness (see cartoon in Figure 4C).

### Generation of SD maps from images series

Time series of images were obtained from TIRF movie in selected regions of atrial myocytes expressing h-K_V_1.5-GFP. SD images were constructed from image stacks containing 30 images corresponding to a time series of 600 seconds using the Z-projection/Standard-Deviation plugin of ImageJ^28^. The standard deviation (SD) map representing the variation in fluorescence intensity for each pixel location in the raw images was used to highlight the h-K_V_1.5-GFP mobility and was overlapped with average fluorescence images (fixed: Z-projection/Average) of microtubules or actin cytoskeleton from the same time series.

### Transmission electron microscopy

For thin section, pieces of atria (1mm^3^) were fixed for 4h in 2% glutaraldehyde in 0.2 M cacodylate buffer, pH=7.2 at RT, then post-fixed with 1% osmium tetroxide containing 1.5% potassium cyanoferrate and contrasted with uranyl acetate 2% in water. After a gradual dehydration in ethanol (solutions from 30% to 100%), the samples were embedded in Epon (Delta microscopie – Labège France). Thin sections of 70 nm were collected onto 200 mesh cooper grids, and counter stained with lead citrate before observation with Hitachi HT7700 electron microscope operated at 80 kV (MIMA2-UMR1131 GABI, Plateau de Microscopie Electronique, Jouy-en-Josas, France). Images were acquired with a charge-coupled device camera AMT–Hitachi (Elexience, Verrière le Buisson, France).

### Image analysis

Fluorescence quantification and colocalization analysis were performed using ImageJ software (NIH) or Imaris 8 and Imaris 9 softwares (Bitplane). Stacks of 4 µm-thick were used for perspectives views/volume rendering, otherwise 3 consecutive stacks were used for image analysis (z-projections/average mean intensity) using ImageJ.

### Human samples

In accordance with approval of Institution Ethic Committee, biopsies of right atrial appendages were obtained from 15 patients undergoing cardiac surgery. Two groups of patients were defined. The control group was constituted of patients in sinus rhythm, preserved left ventricular function and normal atria (n=8); the diseased group was constituted of patients with permanent AF and undergoing cardiac surgery for valve replacement (n=7-8).

### Statistics

Data were tested for normality using the D’Agostino and Pearson normality test. Statistical analysis was performed using Student’s *t* test or ANOVA followed by post-hoc Student-Newman-Keuls test on raw data. Results are given as means ± SEM and *P* values of less than 0.05 were considered significant (* P<0.05, ** P<0.01 and *** P<0.001; ns: not significant). N correspond to the number of experimental replicates, and n to the number of independent biological samples.

## Supporting information

Supporting Informations

## ACKNOWLEDGMENTS

We thank Dr. C. Longin, MIMA2-UMR1131 GABI, Plateau de Microscopie Electronique, Jouy-en-Josas, France for her help with electron microscopy acquisitions.

