## Supporting Informations for "Microtubule polymerization state and clathrin-dependent internalization regulate dynamics of cardiac potassium channel"

**SUPPORTING INFORMATION**

**SUPPLEMENTAL FIGURE LEGENDS**

**Table 1: Summary of h-$K_V$1.5-EGFP channel dynamics parameters in the membrane plane of atrial myocytes.** All parameters have been compared to the corresponding control condition using one-way ANOVA. Data are expressed as mean±SEM. ns, not significant; * P<0.05; ** P<0.01; *** P<0.001. n=10-15 cells/condition. Average track analyzed/movie=1-7. Note that some conditions (for example colchicine) dramatically reduced the number of trackable particles.

**Supplemental Figure S1. Clathrin localization in atrial myocytes. A,** High resolution 3-D deconvolution microscopy images of clathrin LC in freshly isolated myocytes, and enlargement images of intercalated disc (ID) and lateral membrane (LM). **B,** Transmission electron microscopy images of clathrin vesicles in atrial myocardium at the level of ID, LM and z-line. **C,** Cartoon depicting two different structures associated with clathrin: clathrin buds (bud) and clathrin-coated vesicles (CCV). Arrow heads in **B** point to buds; arrows point CCV. **D,** Quantification of buds *vs* CCV at the LM and ID from transmission electron microscopy images.

**Supplemental Figure S2. Effect of clathrin blockade on $K_V$1.5 anterograde trafficking. A,** Trafficking block and release assay: atrial myocytes overexpressing h-$K_V$1.5-HA were treated with brefeldin A to prevent ER/Golgi transport and trafficking was restarted for 2 h before addition of sucrose (or control). (Left) Example images acquired in high resolution 3-D deconvolution microscopy of cells live-stained with anti-HA (surface HA, green), fixed/permeabilized and post-stained with anti-HA (total HA, red). (Right) Quantification of surface HA staining relative to total HA. ** P<0.01; n=number of cells. **B,** High resolution 3-D deconvolution image of clathrin light chain staining in freshly isolated adult rat atrial myocyte (and enlargement below) showing the high concentration of clathrin vesicles surrounding the nucleus in compartments corresponding to the Golgi network. **C,** Transmission electron microscopy images taken from adult rat atria at different enlargements. The Golgi network is highlighted in green, clathrin vesicles in pink.

**Supplemental Figure S3. Activation properties of $I_{Kur}$ current upon clathrin blockade. A,** Calculation of the activation-voltage relationships of potassium current-density before and after sucrose treatment showed no significant change ($V_{0.5}$ for control and sucrose treatment were respectively 6.8 ± 3.3 mV (n=9) and 10.8 ± 2.1 mV (n = 11) as for corresponding slope factors k which were 14.1 ± 1.9 mV and 15.2 ± 1.4 mV). **B,** Calculation of the activation-voltage relationships of potassium current-density in control scrambled siRNA and si-clathrin light chain RNA showed no change ($V_{0.5}$ for si-scramble and si-clathrin treatments were respectively 16.4 ± 4.8 mV (n=8) and 19.8 ± 4.7 mV (n = 7) with corresponding slope factors k of 14.2 ± 1.9 mV and 16.4 ± 2.3 mV). Activations curves clearly demonstrate that there is no shift in the activation properties of $K_V$1.5-mediated potassium current upon clathrin blockade.

**Supplemental Figure S4. Example images showing that sucrose treatment does not affect neither cell morphology nor GFP distribution. A,** Example differential interference contrast (DIC) images taken before sucrose treatment (left) and after sucrose treatment (right). **B,** GFP alone (Ad-GFP) has a diffuse distribution in atrial myocytes and is not affected by sucrose treatment (final minutes: 2 hours treatment).

**Supplemental Figure S5. Effect of cytoskeleton disrupting agents on $K_V$1.5 surface expression. A,** Current density-voltage relationships of endogenous $I_{Kur}$ obtained from atrial myocytes in control (EtOH) and colchicine conditions and representative traces in the insert (holding potential: -80 mV; test potential: +60 mV). **B,** Current density-voltage relationships of endogenous $I_{Kur}$ obtained from atrial myocytes in control (DMSO) and cytochalasin D conditions and representative traces in the insert (holding potential: -80 mV; test potential: +60 mV). **C,** Example images of atrial myocytes transduced with h-$K_V$1.5-HA and live-stained with anti-HA after treatment with colchicine or cytochalasin D (and respective controls). **D,** Summary graphs of surface HA staining after treatment with disrupting agents. ns, not significant; n=number of cells.

**Supplemental Figure S6. Effect of cytoskeleton disrupting agents on h-$K_V$1.5-HA anterograde trafficking and internalization. A,** Anterograde trafficking: atrial myocytes overexpressing h-$K_V$1.5-HA were treated with brefeldin A to prevent ER/Golgi transport and trafficking was restarted for 2 h before addition of cholchicine or cytochalasin D (or respective controls EtOH and DMSO). (Left) Example images acquired in high resolution 3-D deconvolution microscopy of cells live-stained with anti-HA (surface HA, green), fixed and post-stained with anti-HA (total HA, red). (Right) Quantifications of surface HA staining relative to total HA. **B,** Retrograde trafficking: after ER/Golgi transport block, cells were treated with cytoskeleton disrupting agents (or respective controls) and live-stained with anti-HA antibody. Some cells were immediately fixed (T=0) or returned back to the incubator for 2 hours before fixation (T=2h). (Left) Example images of cells acquired in high resolution 3-D deconvolution microscopy at the two time-points. (Right) Summary graphs of the fluorescent HA staining measured at cell boundaries using corresponding DIC images. * $P<0.05$; ** $P<0.01$; *** $P < 0.001$; n=number of cells.

**Supplemental Figure S7. Effect of cytoskeleton disrupting agents on h-$K_V$1.5-HA internalization.** Atrial myocytes overexpressing h-$K_V$1.5-HA were treated with brefeldin A for 16 hours to prevent ER/Golgi transport. Cells were then treated for 2 h with EtOH or colchicine (microtubule disruption), or for 24 h with DMSO or cytochalasin D (actin cytoskeleton disruption) before been live stained with rabbit anti-HA antibody and goat anti-rabbit Fab A488 antibody. **A)** Example images of cells immediately fixed after live staining (T = 0). **B)** example images of cells incubated for two additional hours (T=2 h, internalization assay). Top images correspond to the entire z-projection of 20 stacks acquired at 0.2 µm intervals in the z axis (grid: 5X5 µm). Cell surface perimeters have been delineated manually and surface rendering shown in grey has been done using Imaris' surface module. Bottom images are perspective views of the corresponding 3-D-rendered images. Note that the HA staining is restricted to cell boundaries at T=0 and that channels moving inside the cell can be observed after 2 h internalization.

#### SUPPLEMENTAL MOVIE LEGENDS

**Movie 1. Dynamics of h-$K_V$1.5-GFP vesicles in atrial myocytes and effect of clathrin blockade.** Adult rat atrial myocytes were transduced with h-$K_V$1.5-GFP. Images were analyzed by time-lapse TIRF microscopy with an angle of illumination for maximum penetration of the evanescent field into the cell enabling an imaging depth of 70 nm using an Olympus IX81-ZDC2 microscope and the Cell$^{tirf}$ system. **(A, C)** In a first set of experiments frames were acquired every 20 seconds for 2 hours in control **(A)** or in the presence of 225 mmol/L sucrose **(C)**. **(B, D)** In order to better evaluate the fast dynamics of h-$K_v$1.5-GFP particles, in a second set of experiments frames were taken every 0.5 seconds over a period of 2 minutes in control **(B)** or in the presence of 225 mmol/L sucrose **(D)**. Both sets of experiments are represented in the accelerated video. Moving particles and large sucrose clusters are tracked in red.

**Movie 2. Effect of microtubule depolimerization on h-$K_V$1.5-GFP vesicle dynamics in atrial myocytes.** Adult rat atrial myocytes were transduced with h-$K_V$1.5-GFP. Images were analyzed by time-lapse TIRF microscopy with an angle of illumination for maximum penetration of the evanescent field into the cell enabling an imaging depth of 70 nm using an Olympus IX81-ZDC2 microscope and the Cell$^{tirf}$ system. Myocytes expressing h-$K_V$1.5-GFP were incubated with 10 µmol/L colchicine and frames were acquired every 20 seconds for 2 hours

**(A)** or every 0.5 seconds for 2 minutes **(B)** and are presented in the accelerated video. Moving particles are tracked in red, immobile clusters are tracked in blue.

**Movie 3. Effect of actin depolimerization on h-$K_V$1.5-GFP vesicles in atrial myocytes.** Adult rat atrial myocytes were transduced with h-$K_V$1.5-GFP. Images were analyzed by time-lapse TIRF microscopy with an angle of illumination for maximum penetration of the evanescent field into the cell enabling an imaging depth of 70 nm using an Olympus IX81-ZDC2 microscope and the Cell$^{tirf}$ system. Myocytes expressing h-$K_V$1.5-GFP were incubated with 10 µmol/L cytochalasin-D and frames were acquired every 5 seconds for 12.5 minutes **(A)** or every 0.5 seconds for 2 minutes **(B)** and are presented in the accelerated video. Moving particles are tracked in red.

### Supplemental Figure S1

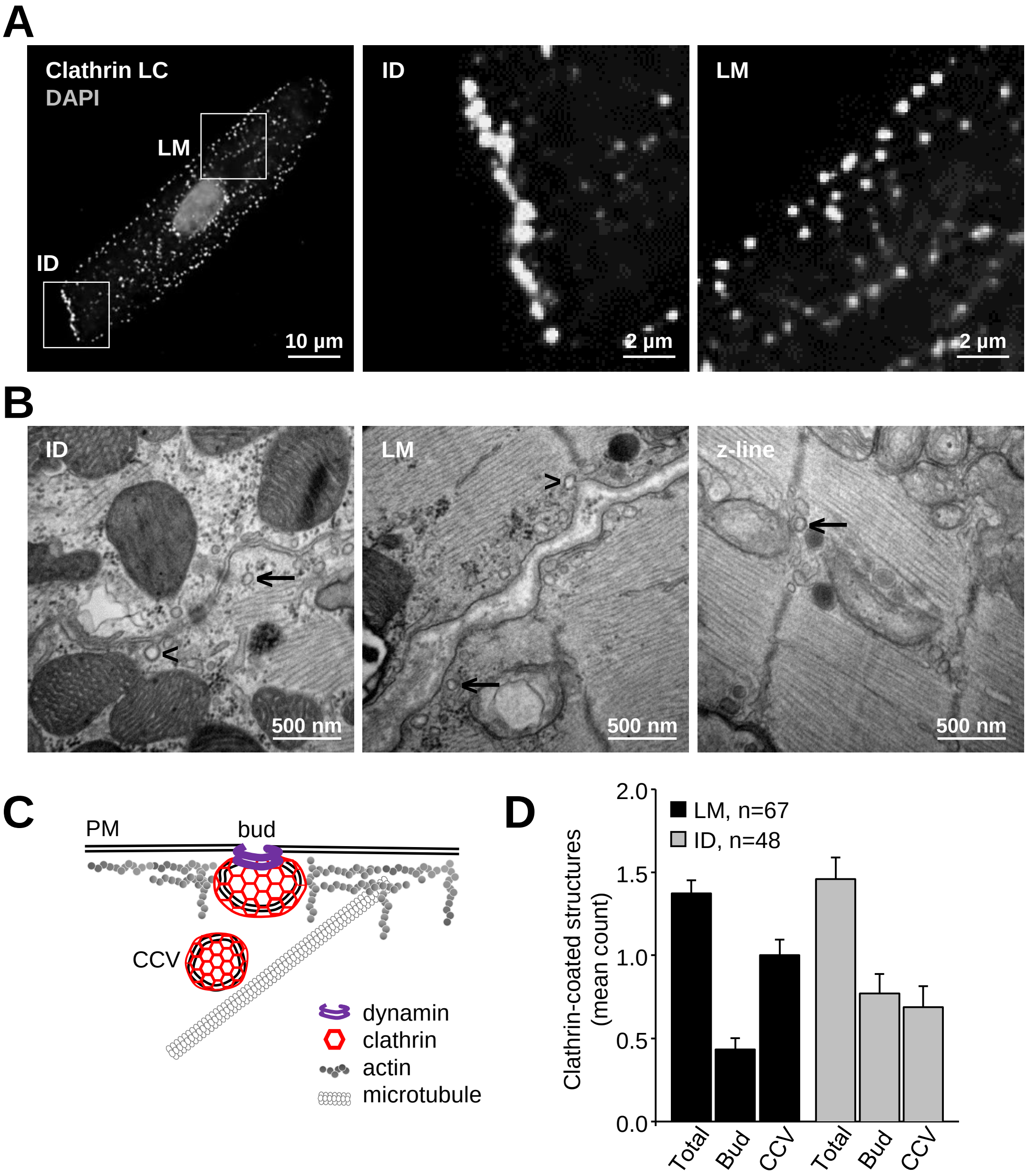


**Figure S1. Clathrin localization in atrial myocytes. A,** High resolution 3-D deconvolution microscopy images of clathrin LC in freshly isolated myocytes, and enlargement images of intercalated disc (ID) and lateral membrane (LM). **B,** Transmission electron microscopy images of clathrin vesicles in atrial myocardium at the level of ID, LM and z-line. **C,** Cartoon depicting two different structures associated with clathrin: clathrin buds (bud) and clathrin-coated vesicles (CCV). Arrow heads in **B** point to buds; arrows point CCV. **D,** Quantification of buds *vs* CCV at the LM and ID from transmission electron microscopy images.

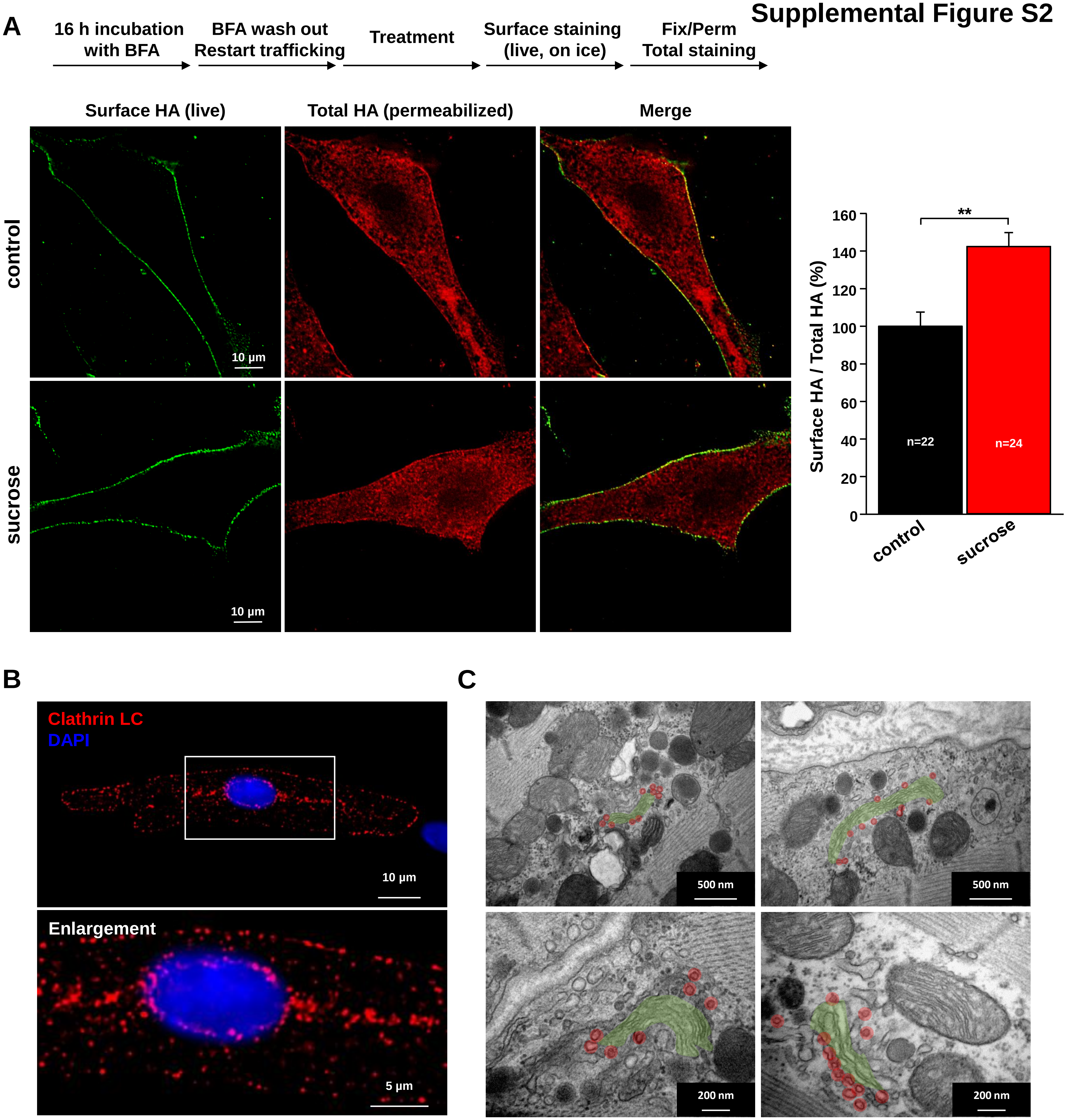


**Supplemental Figure S2. Effect of clathrin blockade on $K_V$1.5 anterograde trafficking. A,** Trafficking block and release assay: atrial myocytes overexpressing h-$K_V$1.5-HA were treated with brefeldin A to prevent ER/Golgi transport and trafficking was restarted for 2 h before addition of sucrose (or control). (Left) Example images acquired in high resolution 3-D deconvolution microscopy of cells live-stained with anti-HA (surface HA, green), fixed/permeabilized and post-stained with anti-HA (total HA, red). (Right) Quantification of surface HA staining relative to total HA. ** P<0.01; n=number of cells. **B,** High resolution 3-D deconvolution image of clathrin light chain staining in freshly isolated adult rat atrial myocyte (and enlargement below) showing the high concentration of clathrin vesicles surrounding the nucleus in compartments corresponding to the Golgi network. **C,** Transmission electron microscopy images taken from adult rat atria at different enlargements. The Golgi network is highlighted in green, clathrin vesicles in pink.

### Supplemental Figure S3

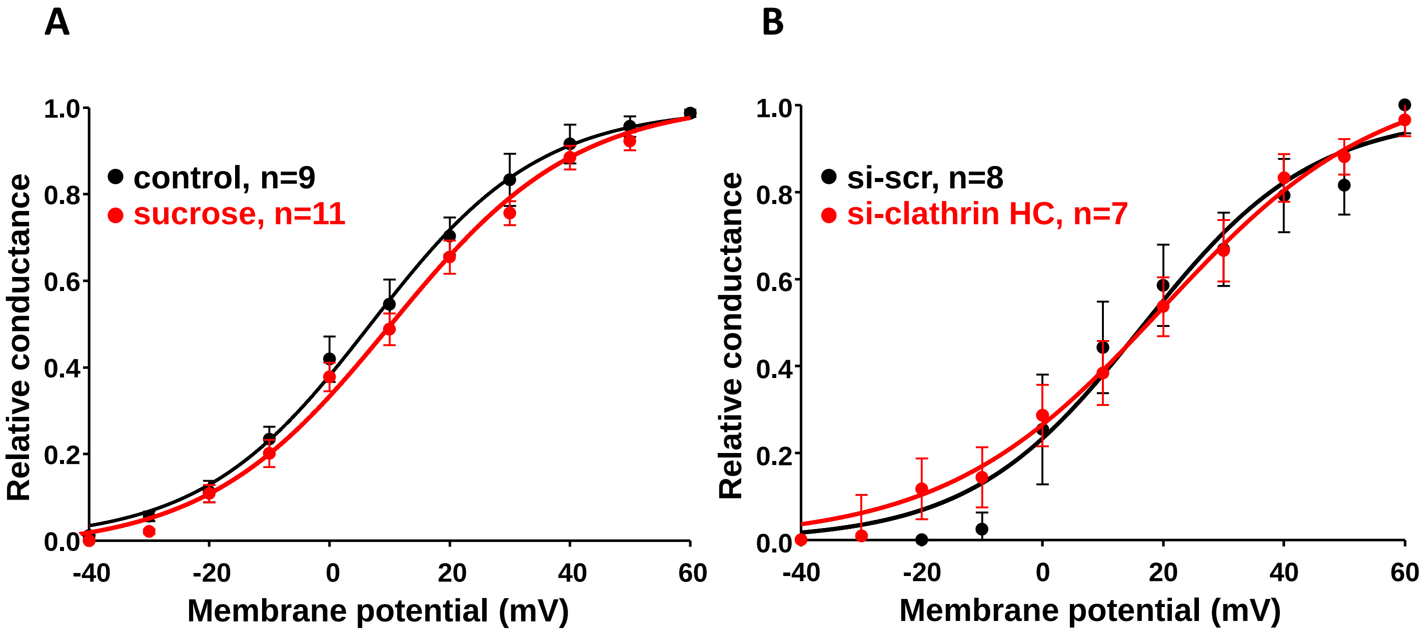


**Supplemental Figure S3. Activation properties of $I_{Kur}$ current upon clathrin blockade. A,** Calculation of the activation-voltage relationships of potassium current-density before and after sucrose treatment showed no significant change ($V_{0.5}$ for control and sucrose treatment were respectively 6.8 ± 3.3 mV (n=9) and 10.8 ± 2.1 mV (n = 11) as for corresponding slope factors k which were 14.1 ± 1.9 mV and 15.2 ± 1.4 mV). **B,** Calculation of the activation-voltage relationships of potassium current-density in control scrambled siRNA and si-clathrin light chain RNA showed no change ($V_{0.5}$ for si-scramble and si-clathrin treatments were respectively 16.4 ± 4.8 mV (n=8) and 19.8 ± 4.7 mV (n = 7) with corresponding slope factors k of 14.2 ± 1.9 mV and 16.4 ± 2.3 mV). Activations curves clearly demonstrate that there is no shift in the activation properties of $K_V1.5$-mediated potassium current upon clathrin blockade.

### Supplemental Figure S4

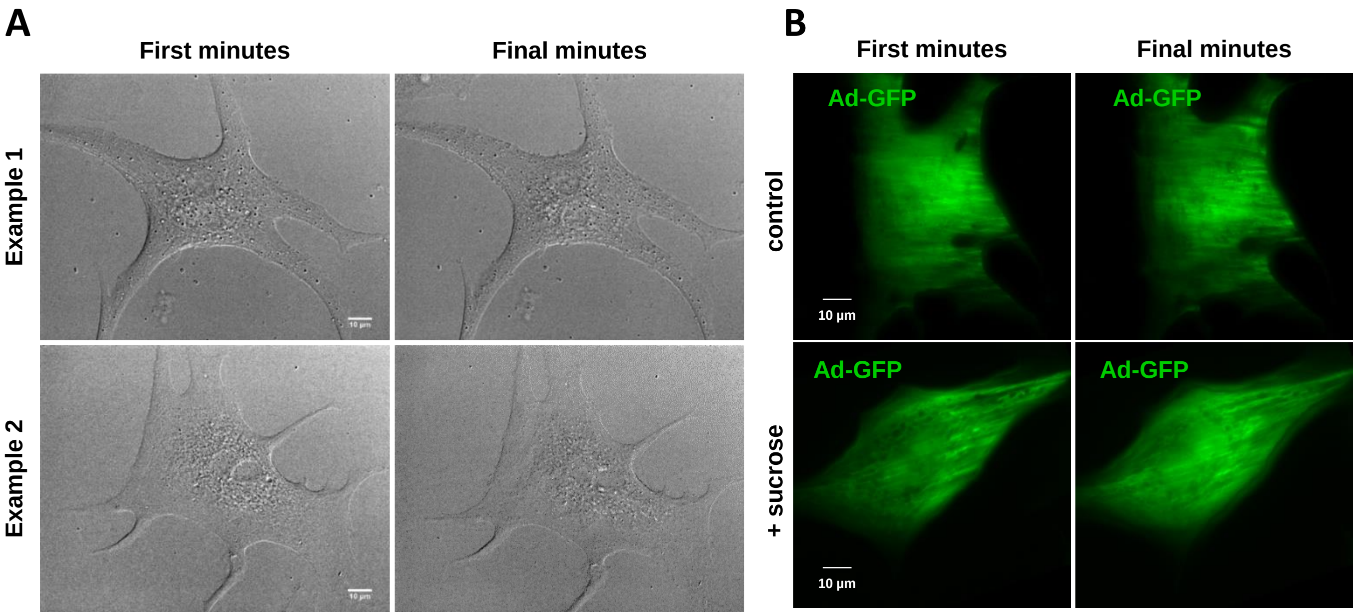


**Supplemental Figure S4. Example images showing that sucrose treatment does not affect neither cell morphology nor GFP distribution. A,** Example differential interference contrast (DIC) images taken before sucrose treatment (left) and after sucrose treatment (right). **B,** GFP alone (Ad-GFP) has a diffuse distribution in atrial myocytes and is not affected by sucrose treatment (final minutes: 2 hours treatment).

Supplemental Figure S5

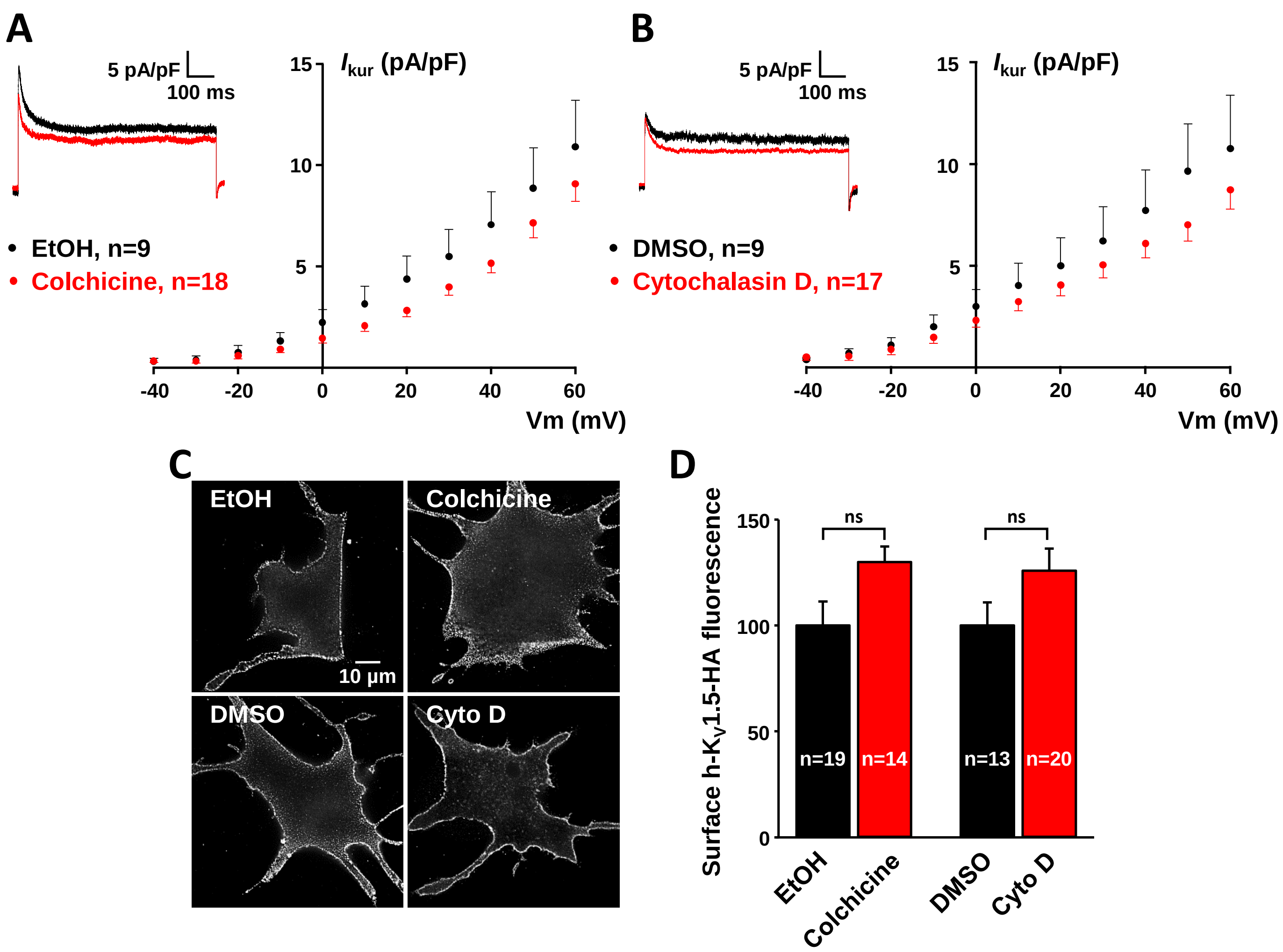


**Supplemental Figure S5. Effect of cytoskeleton disrupting agents on $K_V$1.5 surface expression. A,** Current density-voltage relationships of endogenous $I_{Kur}$ obtained from atrial myocytes in control (EtOH) and colchicine conditions and representative traces in the insert (holding potential: -80 mV; test potential: +60 mV). **B,** Current density-voltage relationships of endogenous $I_{Kur}$ obtained from atrial myocytes in control (DMSO) and cytochalasin D conditions and representative traces in the insert (holding potential: -80 mV; test potential: +60 mV). **C,** Example images of atrial myocytes transduced with h-$K_V$1.5-HA and live-stained with anti-HA after treatment with colchicine or cytochalasin D (and respective controls). **D,** Summary graphs of surface HA staining after treatment with disrupting agents. ns, not significant; n=number of cells.

#### Supplemental Figure S6

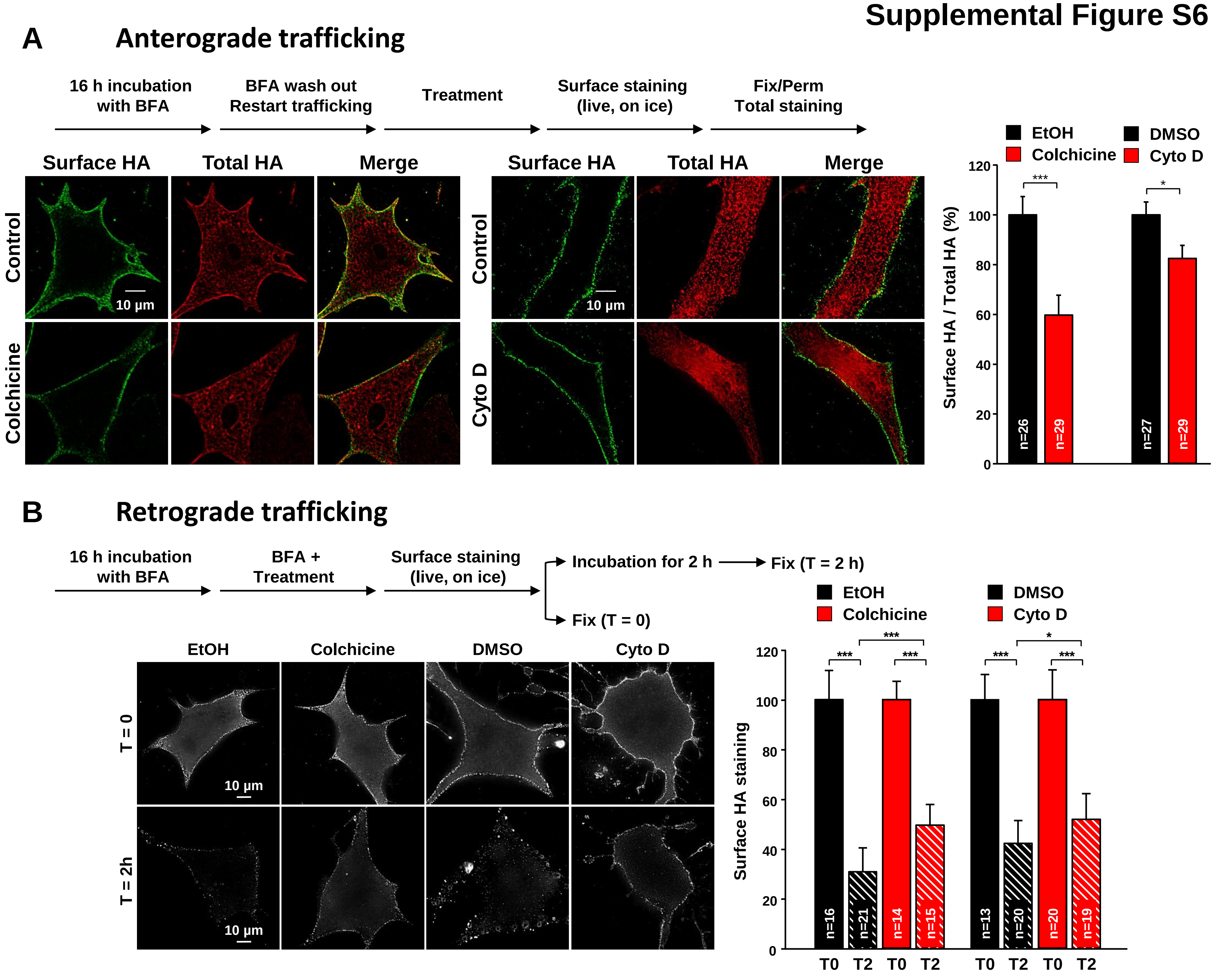


**Supplemental Figure S6. Effect of cytoskeleton disrupting agents on h-$K_V$1.5-HA anterograde trafficking and internalization. A,** Anterograde trafficking: atrial myocytes overexpressing h-$K_V$1.5-HA were treated with brefeldin A to prevent ER/Golgi transport and trafficking was restarted for 2 h before addition of cholchicine or cytochalasin D (or respective controls EtOH and DMSO). (Left) Example images acquired in high resolution 3-D deconvolution microscopy of cells live-stained with anti-HA (surface HA, green), fixed and post-stained with anti-HA (total HA, red). (Right) Quantifications of surface HA staining relative to total HA. **B,** Retrograde trafficking: after ER/Golgi transport block, cells were treated with cytoskeleton disrupting agents (or respective controls) and live-stained with anti-HA antibody. Some cells were immediately fixed (T=0) or returned back to the incubator for 2 hours before fixation (T=2h). (Left) Example images of cells acquired in high resolution 3-D deconvolution microscopy at the two time-points. (Right) Summary graphs of the fluorescent HA staining measured at cell boundaries using corresponding DIC images. * P<0.05; ** P<0.01; *** P < 0.001; n=number of cells.

### Supplemental Figure S7

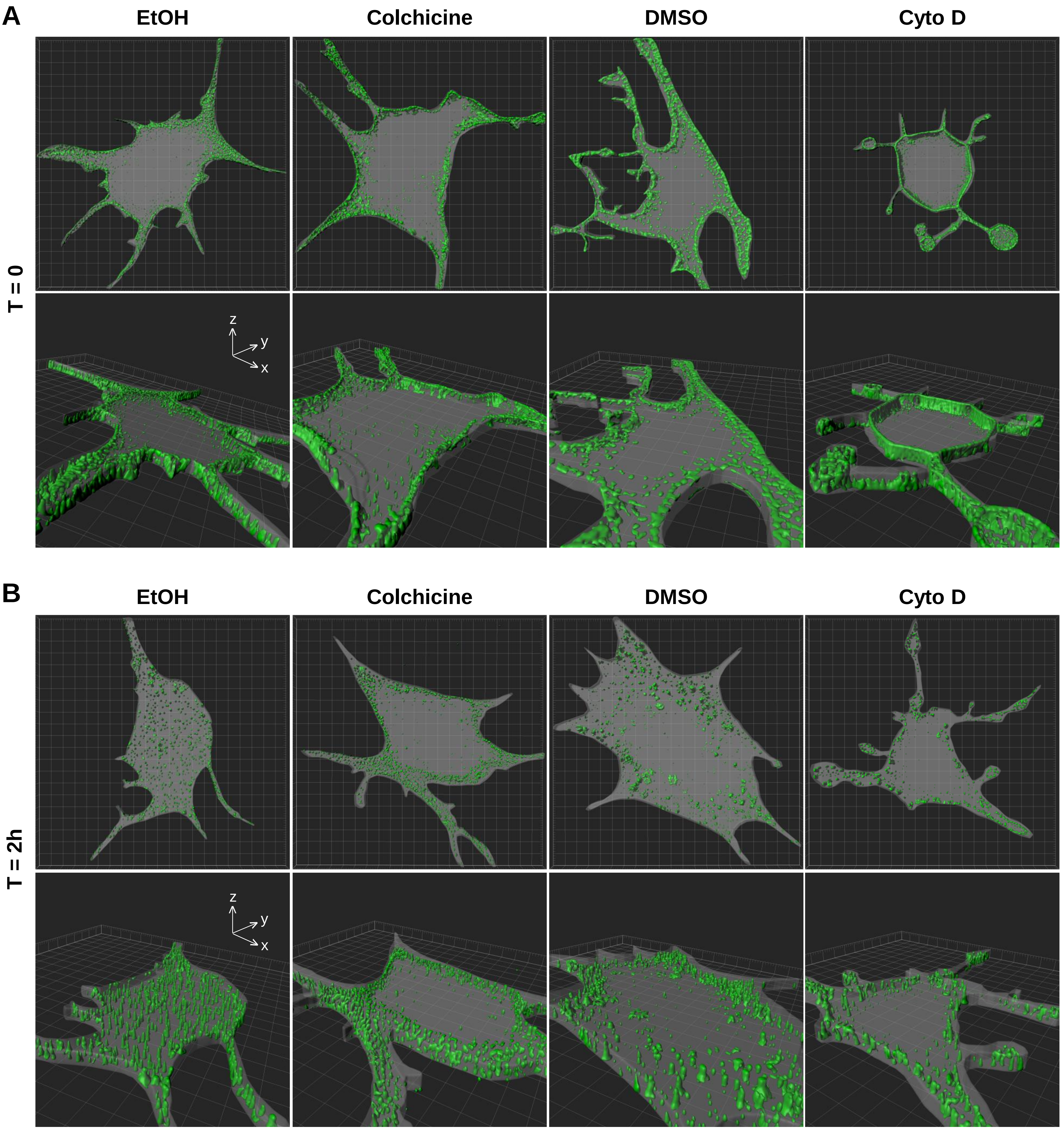


**Supplemental Figure S7. Effect of cytoskeleton disrupting agents on h-$K_V$1.5-HA internalization.** Atrial myocytes overexpressing h-$K_V$1.5-HA were treated with brefeldin A for 16 hours to prevent ER/Golgi transport. Cells were then treated for 2 h with EtOH or colchicine (microtubule disruption), or for 24 h with DMSO or cytochalasin D (actin cytoskeleton disruption) before been live stained with rabbit anti-HA antibody and goat anti-rabbit Fab A488 antibody. **A)** Example images of cells immediately fixed after live staining (T = 0). **B)** example images of cells incubated for two additional hours (T = 2 h, internalization assay). Top images correspond to the entire z-projection of 20 stacks acquired at 0.2 µm intervals in the z axis (grid: 5X5 µm). Cell surface perimeters have been delineated manually and surface rendering shown in grey has been done using Imaris' surface module. Bottom images are perspective views of the corresponding 3D-rendered images. Note that the HA staining is restricted to cell boundaries at T = 0 and that channels moving inside the cell can be observed after 2 h internalization.

### Supplemental Table 1

| | Tracks speed max (µm/s) | Tracks length (µm) | Straightness (0 – 1) |
|---|---|---|---|
| **Control** | 1.05±0.04 | 3.89±0.34 | 0.65±0.03 |
| **Sucrose vesicles** | 1.17±0.06$^{ns}$ | 5.08±0.41* | 0.53±0.03** |
| **Sucrose clusters** | 0.531±0.05*** | 1.19±0.10*** | 0.11±0.01*** |
| **Control-EtOH** | 1.36±0.08 | 4.17±0.37 | 0.63±0.03 |
| **Colchicine** | 0.60±0.10** | 0.10±0.21$^{*}$ | 0.33±0.15$^{*}$ |
| **Control-DMSO** | 1.18±0.06 | 3.10±0.41 | 0.66±0.03 |
| **Cytochalasin D** | 1.27±0.15$^{ns}$ | 3.24±0.38$^{ns}$ | 0.68±0.04$^{ns}$ |

**Supplementary Table 1: Summary of h-$K_V$1.5-EGFP channel dynamics parameters in the membrane plane of atrial myocytes.** All parameters have been compared to the corresponding control condition using one-way ANOVA. Data are expressed as mean±SEM. ns, not significant; * P<0.05; ** P<0.01; *** P<0.001. n=10-15 cells/condition. Average track analyzed/movie=1-7. Note that some conditions (for example colchicine) dramatically reduced the number of trackable particles.
